# Single-Cell Sequencing Reveals Extensive Genetic Diversity Underlying Pediatric ALL Treatment Complexity

**DOI:** 10.1101/2025.03.19.644196

**Authors:** Yakun Pang, Tamara Prieto, Veronica Gonzalez-Pena, Athena Aragon, Yuntao Xia, Sheng Kao, Srinivas Rajagopalan, John Zinno, Jean Quentin, Julien Laval, Dennis J. Yuan, Nathaniel D. Omans, Catherine Potenski, David Klein, Matthew MacKay, Iwijn De Vlaminck, John Easton, William Evans, Dan A. Landau, Charles Gawad

**Author notes:** equal contribution (Yakun Pang, Tamara Prieto). Correspondence: Charles Gawad.

## Abstract

Approximately 15-20% of children with acute lymphoblastic leukemia (pALL) are not cured despite years of treatment with more than ten chemotherapeutic agents, yet bulk sequencing reveals minimal somatic variation that could explain genetic treatment resistance. Using error-corrected bulk sequencing, and primary template-directed amplification (PTA)-enabled single-cell exome and whole-genome sequencing, we discovered much more genetic diversity in pALL. Individual leukemic cells harbor 3.7-fold more mutations (mean 3,553) than detected in bulk samples (mean 965). Many ETV6-RUNX1 patients harbor multiple independent ras-mutated clones, and we also identified convergent evolution of *JAK2* mutations as well as copy number alterations. PTA-based phylogenetic analysis of over 150 single-cell genomes linked heritable phenotypes to specific genetic alterations and identified low-frequency clones harboring mutations in treatment resistance-associated genes (*ITGA9*, *NTRK3*, *SULF2*) that increased in frequency during treatment. These findings reveal extensive hidden genetic diversity in pALL while providing a framework for the early detection of high-risk leukemic populations.

## Introduction

Pediatric acute lymphoblastic leukemia (pALL), the most common childhood cancer, is characterized by two seemingly contradictory observations. Bulk sequencing of pALL tissue samples reveals one of the lowest mutation burdens among all cancers at 0.1-1 mutations per megabase^1,2^ while curative treatment regimens are some of the most complex, requiring 10-15 chemotherapeutic agents over 2-3 years^3^, suggesting there are additional factors that influence treatment responses of specific cellular populations. Moreover, this disconnect between the apparent genetic simplicity and treatment complexity suggests that either genetic diversity has little influence on treatment response or that true genetic diversity is not being measured with bulk sequencing.

Recent single-cell RNA sequencing studies have begun revealing hidden cellular complexity in pediatric leukemia samples by establishing comprehensive atlases of childhood leukemia cells ^4^ and linking developmental states to drug response^5,6^. However, these transcriptional studies are unable to measure the genetic or epigenetic alterations within those populations, and thus cannot explain why genetically “simple” leukemia requires prolonged multi-agent therapy. In addition, pharmacogenomic studies that compare bulk genomic measurements at diagnosis to drug response has identified drug sensitivity patterns across genetic subtypes between patients^7^. However, this strategy is again relying on bulk sequencing before drug exposure, which cannot determine whether there are genetic subpopulations of cells within a patient that exhibit varied drug susceptibility.

Existing evidence suggests that are able to measure genetic changes in ALL samples over time suggest that pALL’s genetic diversity is currently underestimated. For example, bulk sequencing studies have detected multiple co-existing clones at diagnosis^2,8^, and serial analyses have revealed substantial clonal evolution during treatment^9,10^. Traditional flow cytometric measurements of residual disease have detected disease at the end of induction therapy in about 15% of patients, while newer, more sensitive strategies for monitoring minimal residual disease (MRD) have found that 25-30% of patients are MRD positive after four to six weeks of remission induction therapy^11,12^. This suggests that rare subclones with variable treatment sensitivity are present at low levels in a significant fraction of patients. However, little is known about why those rare cells are able to survive, as they cannot be studied with standard bulk sequencing methods.

Here, we employed primary template-directed amplification (PTA) to perform single-cell whole genome sequencing with high accuracy^13^. Using novel computational strategies with these data, combined with error-corrected sequencing that detects mutations at a frequency of 0.1%^14^, we characterized the subclonal genetic landscape of pALL at single-cell resolution **(Figure 1A).** By integrating genetic, surface marker expression, and clinical data in the same cells, we were then able to associate specific genetic changes with cellular phenotypes, including treatment response.

**Figure 1.**
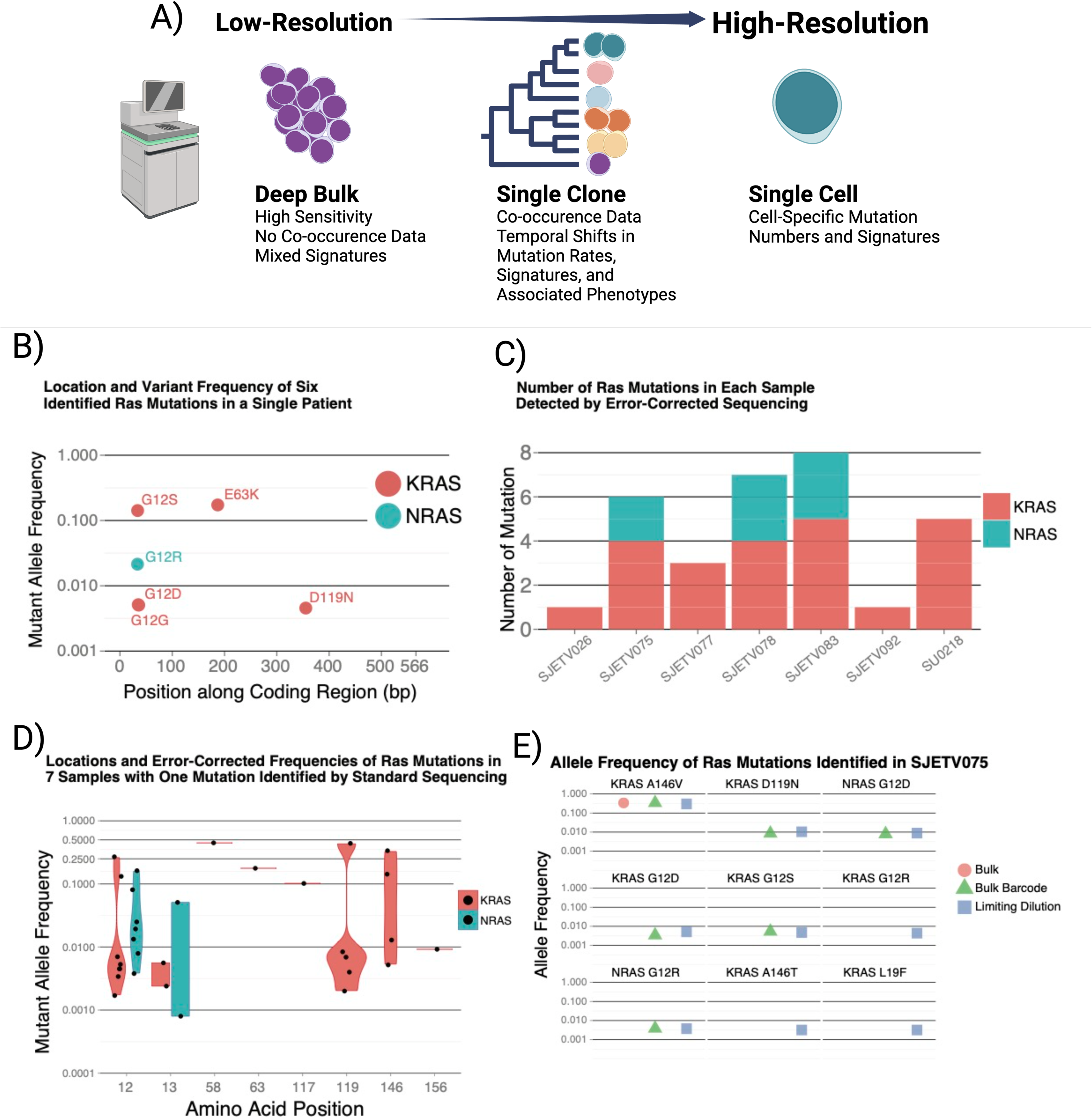
Evidence for large population genetic diversity in *Ras-mutant* pALL. (A) Overview of additional insights we obtained in this study by studying pALL at increasing levels of resolution. (B) Error-corrected sequencing in a patient with 1 ras mutation detected with bulk sequencing identified 4 additional lower-frequency activating mutations. (C) Subclonal ras mutations were also common in a larger cohort in which each patient had one mutation identified in the bulk sample but had a median of 5 activating ras mutations. (C) The allele frequency distributions of ras mutations show no evidence of preferential selection of specific amino acid changes. (D) The increased sensitivity of mutation detection with limiting dilution identifies a total of seven activating ras mutations in a patient that had only one ras mutation detected with bulk sequencing.

With this approach, we discovered that individual pALL cells harbor 3,553 mutations versus 965 detected by bulk sequencing, revealing previously undetectable population-level diversity. Within patients, we found that convergent evolution generates multiple independent ras and JAK2 mutations in distinct clones. In addition, with our phylogenetic tree construction, we order the timing of mutations and mutational signatures while also connecting genotype to phenotype.

Finally, pre-existing rare or undetectable clones with relapse-associated mutations in ITGA9, NTRK3, and SULF2 were detected in surviving leukemia cells after four to six weeks of therapy. Taken together, these findings reshape our view of pALL from a genetically simple to a genetically complex disease, explaining why multi-agent therapy is essential and providing a broader framework for detecting and characterizing treatment-resistant populations before clinical relapse.

## Results

### Hidden ras mutational diversity in pALL revealed by higher resolution bulk sequencing

We performed error-corrected sequencing of 50 genes known to harbor pALL mutational hotspots, starting with an index patient who had a single KRAS E63K mutation identified by bulk sequencing **(Figure S1)**. Error-corrected sequencing revealed four additional lower-frequency KRAS and NRAS mutations, with the NRAS mutation in codon 12, while KRAS mutations were at established activating sites at codons 12, 63, and 119 **(Figure 1B)**. This exclusive occurrence at oncogenic positions despite sequencing the entire gene supports positive selection rather than random mutagenesis events within ras genes^15^.

This finding prompted a systematic analysis of 13 ETV6-RUNX1 patients. Remarkably, error-corrected sequencing identified a median of five activating ras mutations in each of six patients that had previously had a single mutation detected by bulk sequencing, whereas it detected none in seven RAS-negative samples **(Figure 1C)**. The mutations affected KRAS codons 12, 13, 119, and 146 and NRAS codons 12 and 13 **(Figure 1D)**. Of the 33 identified mutations, 31 were C-to-T or C-to-G changes, which contrasts with smoking carcinogen signatures in lung cancer, where 60% are C-to-A substitutions^16^.

To further push detection limits by randomly partitioning mutant clones, we performed error-corrected sequencing on diluted subsamples from patient SJETV075, identifying three additional KRAS mutations (G12R, A146T, L19F) for a total of nine activating variants in that patient **(Figure 1E)**. These findings establish that ETV6-RUNX1 ALL subgroups differ fundamentally in ras pathway dependence, with some patients in whom a high-frequency ras clone is detected harboring extensive ras mutational diversity undetectable with conventional bulk sequencing.

### Single-cell sequencing confirms convergent ras evolution

Having established that multiple ras mutations exist, we next used single-cell exome sequencing with multiple displacement amplification (MDA) to investigate whether they represent nested evolution within single clones or convergent evolution across independent lineages. We sequenced three cells from each of five clones from a sample we had previously performed targeted single-cell sequencing, achieving an average of 82% exome coverage in each cell and identifying 10-29 additional coding mutations per clone **(Figure 2A,B, Figure S2)**^8^.

**Figure 2.**
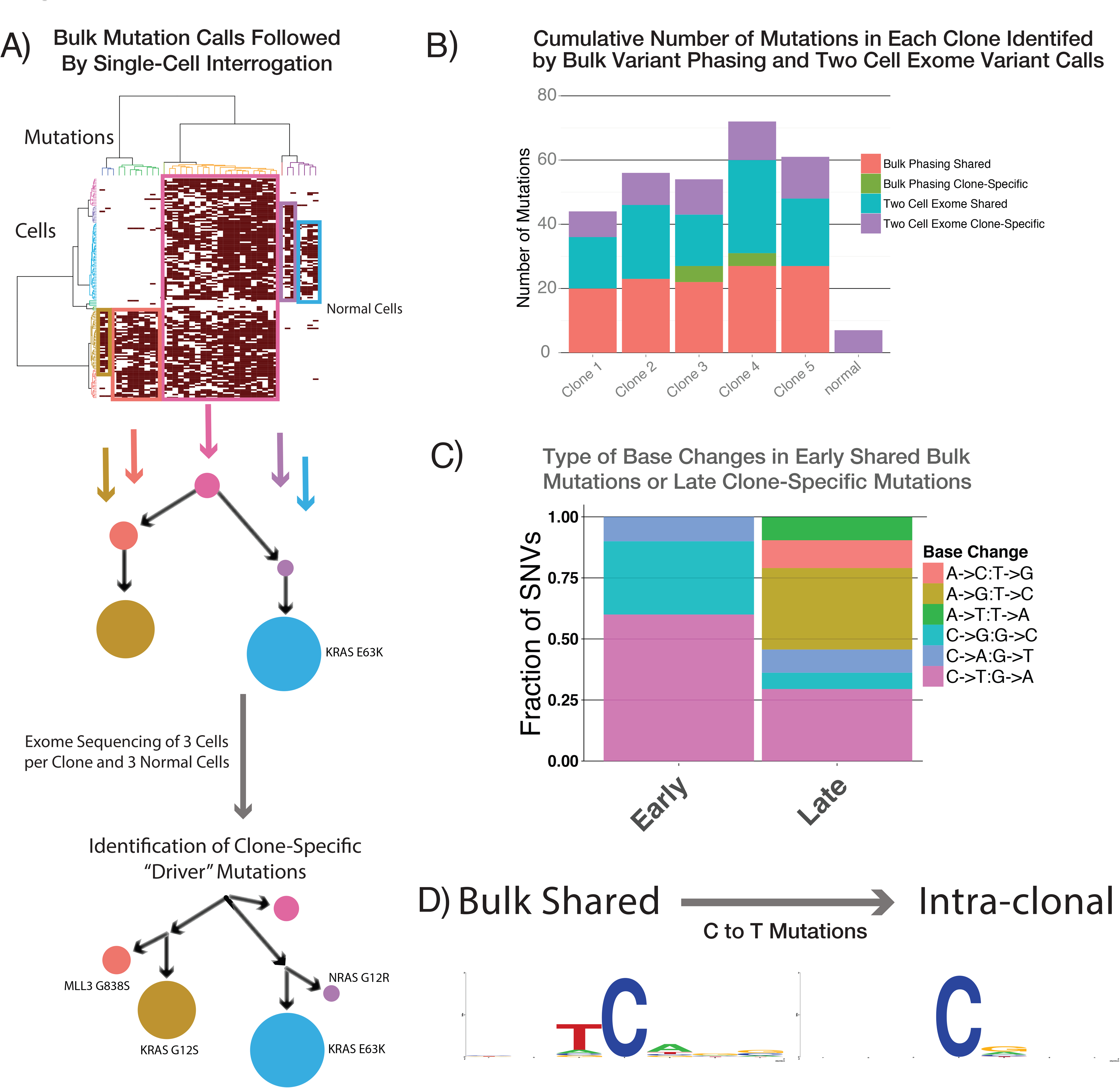
Identification of convergent ras evolution using single-cell exome sequencing. (A) The clonal structure of a diagnostic patient sample that was identified by interrogating single cells for mutations first detected in the bulk sample was further resolved by calling mutations in the single cells alone. The clone-specific *ras* mutations identified as possible causes of the clonal expansions are noted. (B) The number of new mutations identified in each clone using phasing of bulk mutations and 2-cell mutation calls. (C) Base substitution patterns seen in shared (early) and clone-specific (later) mutations. (D) The surrounding motifs in C-to-T mutations in early and late SNVs, showing a the strong APOBEC motif is only present in the early mutations in this patient.

With this approach, in addition to the E63K KRAS clone identified by bulk sequencing, the same ras variants detected in error-corrected sequencing in this patient were also detected. This includes KRAS G12S, which was found to be confined to a minor clone, while NRAS G12D marked a third ras mutant clone, demonstrating that independent lineages convergently evolved ras mutations. This branching architecture reveals selective pressure favoring ras activation across multiple evolutionary trajectories in the same pALL patient. With the limited number of mutations per cell with this approach, we examined potential sources of those mutation. Early mutations shared between clones showed APOBEC signatures (C-to-T and C-to-G at TpC sites), while later clone-specific mutations were predominantly A-to-G and C-to-T without APOBEC motifs **(Figures 2C,D)**.

### PTA-enabled genome sequencing uncovers 3.7-fold higher mutation burden per cell compared to the entire tissue burden detected with bulk sequencing

One limitation of our exome sequencing approach using MDA was that it suffers from severe allelic dropout and imbalances, limiting variant detection. We overcame this barrier using primary template-directed amplification (PTA), achieving >90% variant recall versus 60-70% for MDA **(Figure 3A, S3)**^13^. This technical advance enabled us to measure the actual per-cell mutation burden in pALL cells for the first time where we found a mean of 3,553 mutations per cell, compared to an average of 965 detected in the whole tissues by bulk sequencing **(Figure 3B)**. This represents a 3.7-fold increase in each cell, which occurs across tens to hundreds of billions of cells, resulting in vast population genetic diversity. To further put this mutation burden in context, the 2544 to 4424 mutations per cell are greater than the average number of mutations detected with bulk tissue sequencing across all pediatric tumor subtypes, including those thought to harbor the highest number of mutations, such as osteosarcoma and high-grade gliomas^17^ **(Figure 3C, S4A)**.

**Figure 3.**
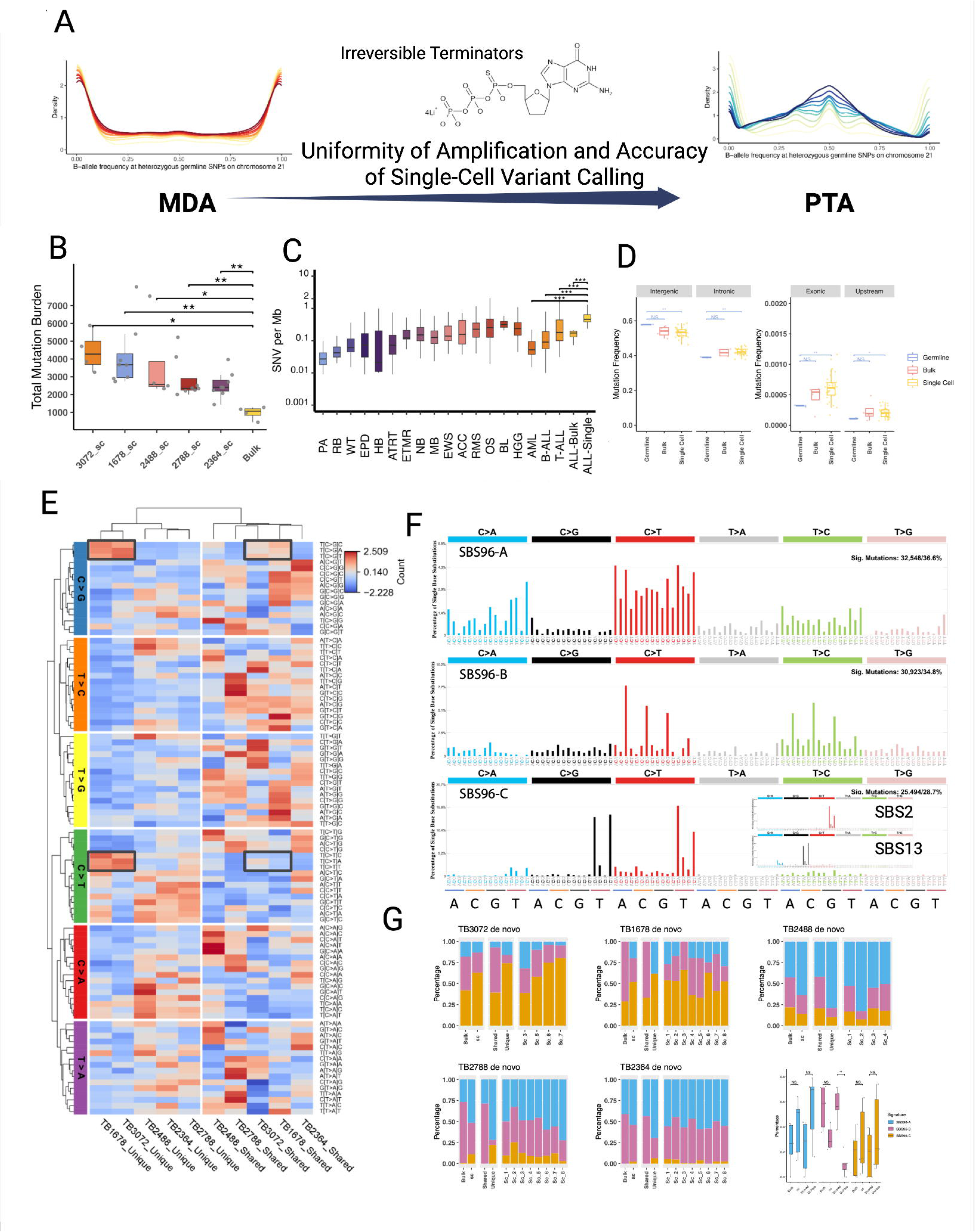
Estimating single-cell mutation rates in ETV6-RUNX1 pALL. (A) Schematic showing improved allelic balance at heterozygous sites enables more accurate small variant calls. (B) Genome-wide mutation numbers in five bulk samples and single cells isolated from the same samples. (C) Comparison of the number of called mutations per megabase in a previously published bulk study of pediatric cancers ^17^, compared to our bulk and single-cell call numbers from five patients. (D) Comparison of germline variant density of specific locations to the fraction of bulk and single-cell variant calls for each of those locations. (E) Heatmap showing the three-base context of unique (late) and shared (early mutations in single cells across five samples. (F) De novo signature detection identified three distinct signatures, with SBS96A (similar to SBS5), SBS96B (similar to SBS1), and SBS96C (similar to APOBEC SBS signatures 2 and 13). (G) The relative abundance of the de novo mutation signatures across patients, which revealed SBS96B consistently decreased over time in these patients.

Further analysis revealed that somatic variants were enriched in intronic and exonic, as well as sequences 1 kb upstream sequences from transcriptional start sites compared to germline variants across the patient’s genome **(Figures 3D, S4B)**. It is unclear if those sites are more prone to mutations at locations with open chromatin, if they are more likely that be changes that can provide a selective advantage for that clone, or if there are other explanations for that pattern. Mutational signature analysis revealed temporal changes: APOBEC signatures (SBS2/SBS13) comprised larger proportions of unique mutations in two patients (1678 and 3072), indicating later APOBEC activity^18^. Conversely, clock-like signatures (SBS1, SBS5) dominated shared mutations, suggesting those were important drivers of mutagenesis early in disease development **(Figure S4C-G)**. We confirmed this finding quantitatively by extracting three *de novo* SBS signatures using Sigprofiler^18^. Signature A resembled COSMIC Signature SBS5, while signature B showed enrichment of C-to-T changes at CpG sites similar to SBS1. Of note, this is a pattern inconsistent with isothermal amplification artifacts, which are de-enriched at CpG sites, supporting these as true variants^19^. Signature C resembled a combination of COSMIC APOBEC signatures SBS2 and SBS13 **(Figure 3F)**. Across samples, the only significant temporal difference was de-enrichment of signature B (similar to SBS1) in later unique mutations. Taken together, these findings revealed that pALL cells harbor far greater mutation burdens than bulk sequencing detects and that mutational signatures dynamically change, with a decrease in spontaneous deamination of CpG sites over time in this cohort **(Figure 3G)**.

### Reproducible single-cell phylogenies validate PTA-mediated variant calling accuracy

Before performing extensive single-cell whole-genome sequencing using PTA, we rigorously validated our capacity to create accurate phylogenetic trees using an *in vitro* evolution model with known lineage relationships. Sequencing 16 cells from related DLD-1 clones, we detected >250,000 somatic SNVs and recovered a phylogeny consistent with our experimental design with maximum bootstrap support **(Figures S6B**, **4A)**.

Critical quality metrics again demonstrated PTA’s superiority compared to MDA with greater preservation of allele frequencies at heterozygous sites and 5.7±4.2% allelic dropout versus 16.4±10.1% for MDA **(Figures S5A,E)**. In addition, PTA achieved 89.4% genome coverage, which is comparable to that of a single-cell colony at the same sequencing depth^20^(**Figure S5B**). Further supporting the validity of our model, branch lengths correlated with culture time (ρ = 0.85, p = 3 × 10⁻⁵) **(Figures 4C)**. Linked-read analysis confirmed 92.4±2.2% somatic variant calling precision and 84.4±7.5% recall **(Figure S5F, S6C)**. Even mutations unique to single cells showed consistent signatures, with <5% unexpected patterns, validating our calling variant accuracy **(Figure 4E)**. We also detected copy-number variants at 5kb resolution (AUC=0.97), enabling integrated genomic analyses **(Figure 4F-G)**.

**Figure 4.**
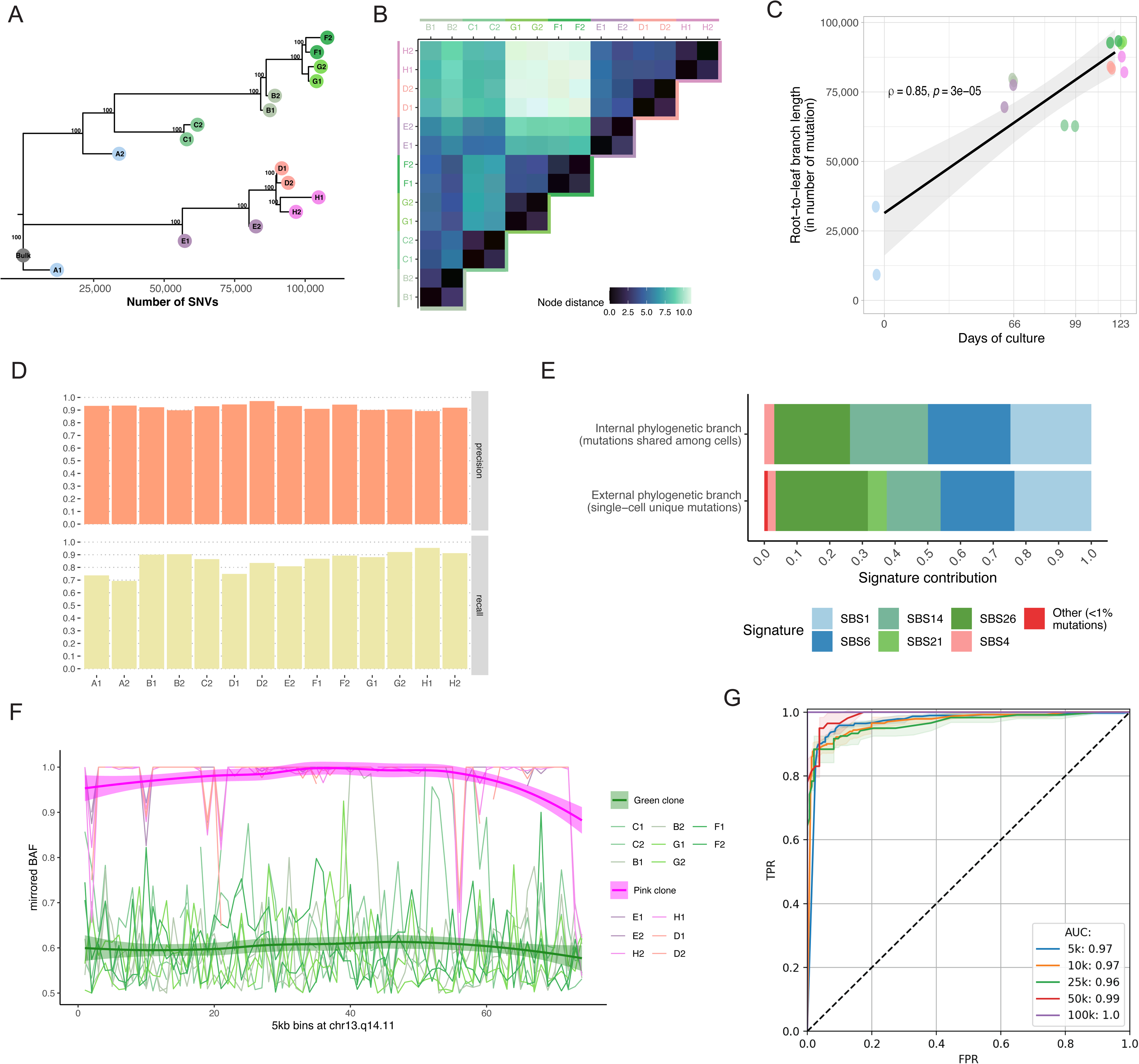
Benchmarking the performance of PTA for scWGS with an *in vitro* evolution experiment (A) Phylogeny of 16 PTA-amplified cells sampled across 7 clones from the evolutionary experiment. Bootstrap values are shown. (C) Node distance of the cells in the reconstructed phylogeny. (D) Correlation between the days of culture of the cells sampled and the branch lengths of the tree in the number of mutations. (E) Recall and precision of the somatic variant calling for the *in vitro* cells. (F) Signature of the mutations mapped to the internal and external phylogenetic branches (G) Mirrored BAF of the in vitro cells (colored as in b) averaged for the heterozygous SNPs contained within 5kb windows which were considered as high-quality for 5 or more cells at a band of the big arm of chromosome 13. (H) ROC curves of the CNV method detection accuracy based on BAF. Cells from the pink clone in b carry a deletion at chr13q14.11, which results in an allelic imbalance (g). Using different BAF fixed thresholds, CNVs were called for every interval across all cells. CNV calls classified as imbalanced in the pink clone cells are considered FNs and those in the green clone are considered FPs. Solid lines show the mean TPR/FPR for 100 groups of cells selected at random with replacement. Lower and upper bounds of the shady areas represent the mean±2*sd of this value.

### Accurate pALL phylogenies enable connecting cellular states with specific genomic alterations

Using this validated tree-building approach, we performed a comprehensive phylogenetic analysis of >150 single-cell genomes from four high-risk pALL patients that had undergone four weeks of induction therapy with three or four chemotherapy drugs but had detectable MRD cells **(Figure 5A)**. All leukemias carried mutations in genes known to be associated with pALL (*JAK2*, *USH2A*, *NRAS*, *ZEB2*, *KRAS*, and *FCGBP*)^2^, enabling classification into premalignant versus malignant cells, which was validated by copy-number patterns and cell surface markers **(Figure 5E,F)**.

**Figure 5.**
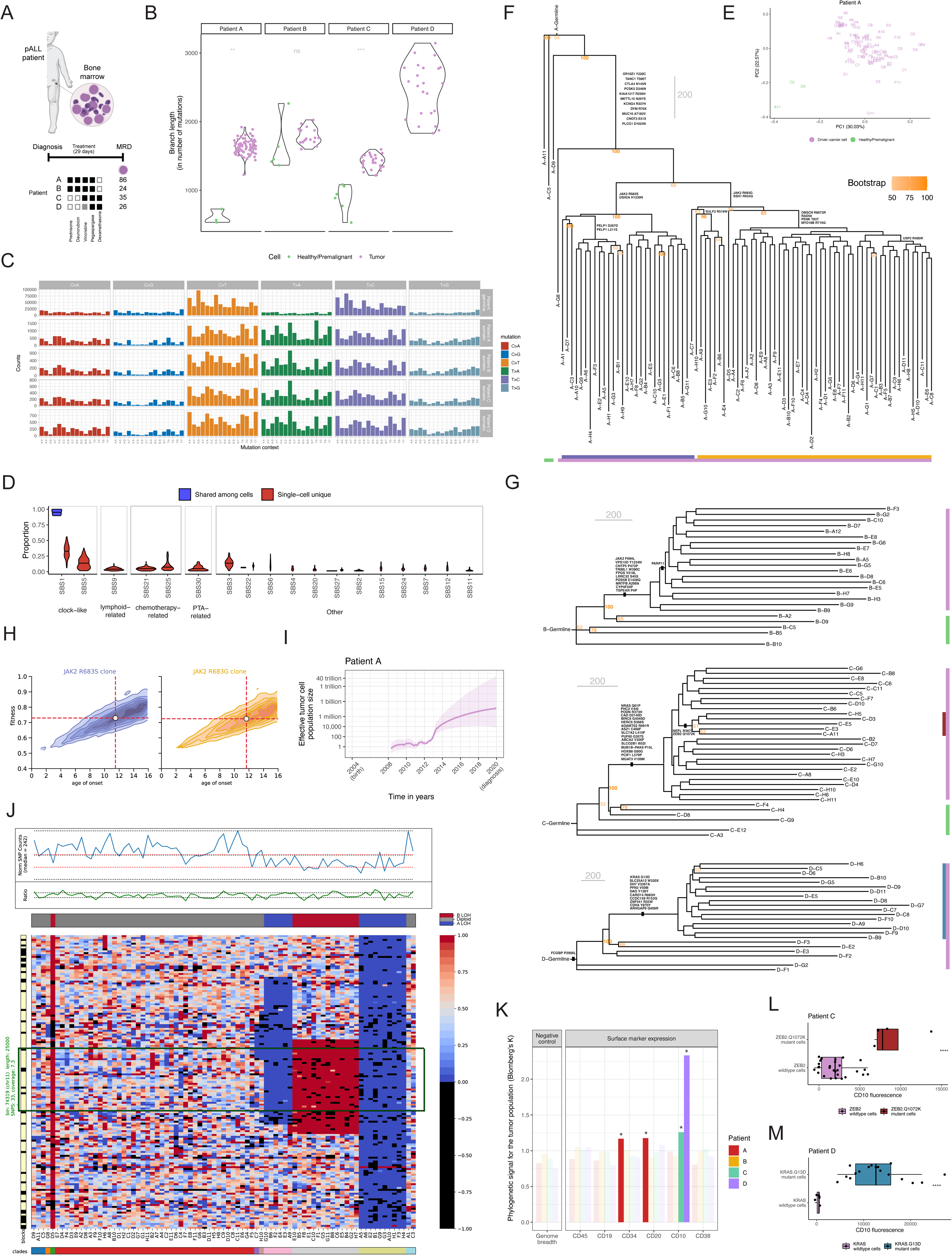
PTA-based scWGS of pALL cells (A) Schematic of the BALL patient, number of cells amplified, and the treatment each patient received. (B) Estimated number of mutations in normal/premalignant and leukemia cells across all four patients (C) Mutational patterns of DLD-1 colorectal cancer compared to 4 pALL patients. (D) Signatures of shared and unique somatic mutations across patients (E) PCA plots of six phenotypic markers across all cells for patient A showing separation of normal/premalignant and leukemia cells (F) Phylogeny of patient A, showing bootstraps of the branches supported in more than 50% of replicates. (G) Phylogenies as annotations for patients B, C, and D. (H) Posterior distribution of the age of onset of the different JAK2 clones using approximate Bayesian computation. (I) Cell population dynamics of tumor cells along lifespan for patients A, B, and C. Solid line represents the posterior median estimate and the shaded region represents the 95% confidence intervals. (J) Mirrored loss-of-heterozygosity (LOH) events in a 25kb window sorted by clade. (K) Phylogenetic signal of the sorting time (negative control) and surface expression for B-cell lineage markers. Non-significant values are shown in lighter colors. (L) Density plots of CD34 and CD20 fluorescence in a clade carrying a 6q deletion compared to all other cells. (M) Violin plots of CD10 fluorescence in cells from patient C with or without a ZEB2 mutations.

In total, we detected ∼130,000 somatic mutations across those 150 cells. Limiting our query to callable regions, malignant cells averaged 1,497±341 (patient A), 1,049±62 (patient B), 1,180±244 (patient C), and 2,253±333 (patient D) somatic SNVs, which exceeded the number of somatic SNVs classified as normal or premalignant B-cells in all patients **(Figure 5B)**. Critically, we detected chemotherapy signatures (SBS21, SBS25)^21^ in variants unique to single cells after just four weeks of treatment, changes that would not be detected until months or years later in patients who relapse **(Figures 5C,D)**.

Phylogenetic analysis showed distinct patterns across patients **(Figures 5E-G)**. We also identified convergent evolution in patient A, whose disease harbored two clones with different JAK2 mutations affecting the same amino acid (R683G and R683S), which also consistently showed similar growth rates by Wright-Fisher modeling with selection in an Approximate Bayesian Computation (ABC) framework **(Figure 5H)**. The presence of both clones before and after treatment can be explained by the fact that they both emerged around the same time interval, as was suggested by the ABC analysis. The similarity in clonal timings and growth rates for both clones was further confirmed by coalescent theory **(Figure 5I)**. Additional analyses detected recurrent copy-number at similar genomic locations that had distinct breakpoints, suggesting convergent evolution of copy number changes in addition to SNVs in that patient. (**Figure 5J**).

Using Blomberg’s K statistic to quantify trait heritability, we discovered significant phylogenetic signals linking genotype to phenotype **(Figure 5K)**. A clade with chromosome 6 deletions (affecting *PRDM1*, *FOXO3*, *HDAC2*) showed increased CD34 and reduced CD20 expression, indicating differentiation arrest at an earlier stage of B-cell development^22^. Similarly, ZEB2-mutant cells expressed higher CD10, connecting specific mutations to the expression of surface markers that are indicative of specific developmental states^23^ **(Figure 5L)**.

### Treatment pressure unmasks pre-existing subclones

The extensive genetic diversity we uncovered suggested that pre-existing mutations in specific clones might influence treatment response. To test this hypothesis, we exposed diagnostic cells from previously sequenced patient SJETV075 to five standard chemotherapy agents (mercaptopurine, vincristine, prednisolone, daunorubicin, and asparaginase) at four concentrations, then tracked clonal dynamics through bulk error-corrected sequencing ^7,24^ **(Figure 6A)**. With our approach, we detected 224 recurrent mutations across all treatment and control conditions, which is five-fold more than standard bulk analysis had detected **(Table S2)**.

**Figure 6.**
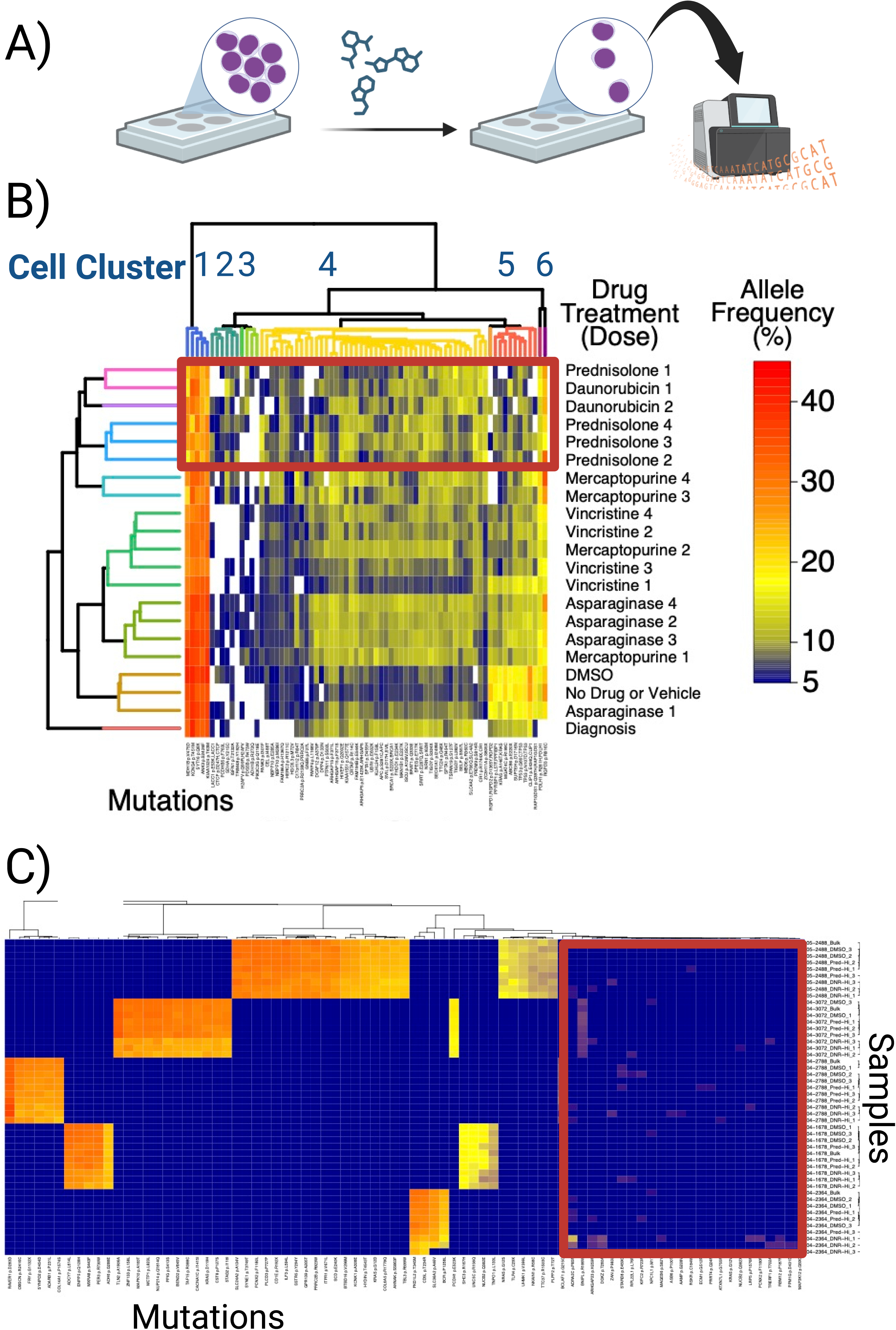
Differential *ex vivo* sensitivity of leukemic populations to chemotherapy. (A) Overview of experimental plan where we treated primary pALL cells with specific drugs, followed by bulk sequencing to identify new variants associated with specific drug treatments. (B) Clusters of mutations showing patterns of response to drug treatment and dosage (increasing dose with higher numbers). Cells with mutations in cluster 6 expanded in culture when compared to the bulk sample, while clones with cluster 5 mutations, which includes *KRAS* A146V and two *TP53* mutations, expanded without treatment or upon exposure to low-dose asparaginase. Mutations in clusters 1 and 4 decreased with increasing doses of vincristine, daunrubicin, prednisolone, and mercaptopurine while clones with mutations in clusters 2 and 3 increased in frequency. All cell died at the highest doses of daunorubicin. (C) Expansion of the concept to five additional patients, where newly detected low-frequency mutations can be detected after exposure to the highest doses of prednisolone or daunorubicin.

Hierarchical clustering revealed patterns of differential response **(Figure 6B)**. The highest-frequency KRAS A146V clone expanded with control treatment but contracted under specific drug combinations. Low-dose mercaptopurine or increasing asparaginase increased the relative frequency of this clone in cluster 5 while prednisolone and daunorubicin showed opposite effects, decreasing clusters 5 while increasing clusters the relative frequency of mutations in cluster 2 to 4. Validation in five additional patients with the highest doses of prednisolone or daunorubicin confirmed patient-specific patterns, where some mutations consistently decreased, while other low-frequency variants became more prevalent **(Figure 6C)**. These findings demonstrate that treatment selection pressure can expose the genetic complexity underlying clonal responses. However, bulk sequencing only provides a list of mutations that cluster together, some of which are difficult to differentiate from background noise. Performing single-cell analyses would provide a less ambiguous, higher resolution view of those drug-resistant clones.

### Resistant clones undetectable at diagnosis can increase in relative frequency during treatment

To determine whether those *ex vivo* observations of emergent clones translate into patients, we performed paired single-cell analyses before and after 4 weeks of combination chemotherapy, when the leukemic cell burden decreases from about 100 billion to 10-100 million cells^25^ **(Figure 7)**. This temporal analysis at single-cell resolution again revealed dynamic changes in the relative frequencies of clonal populations.

**Figure 7.**
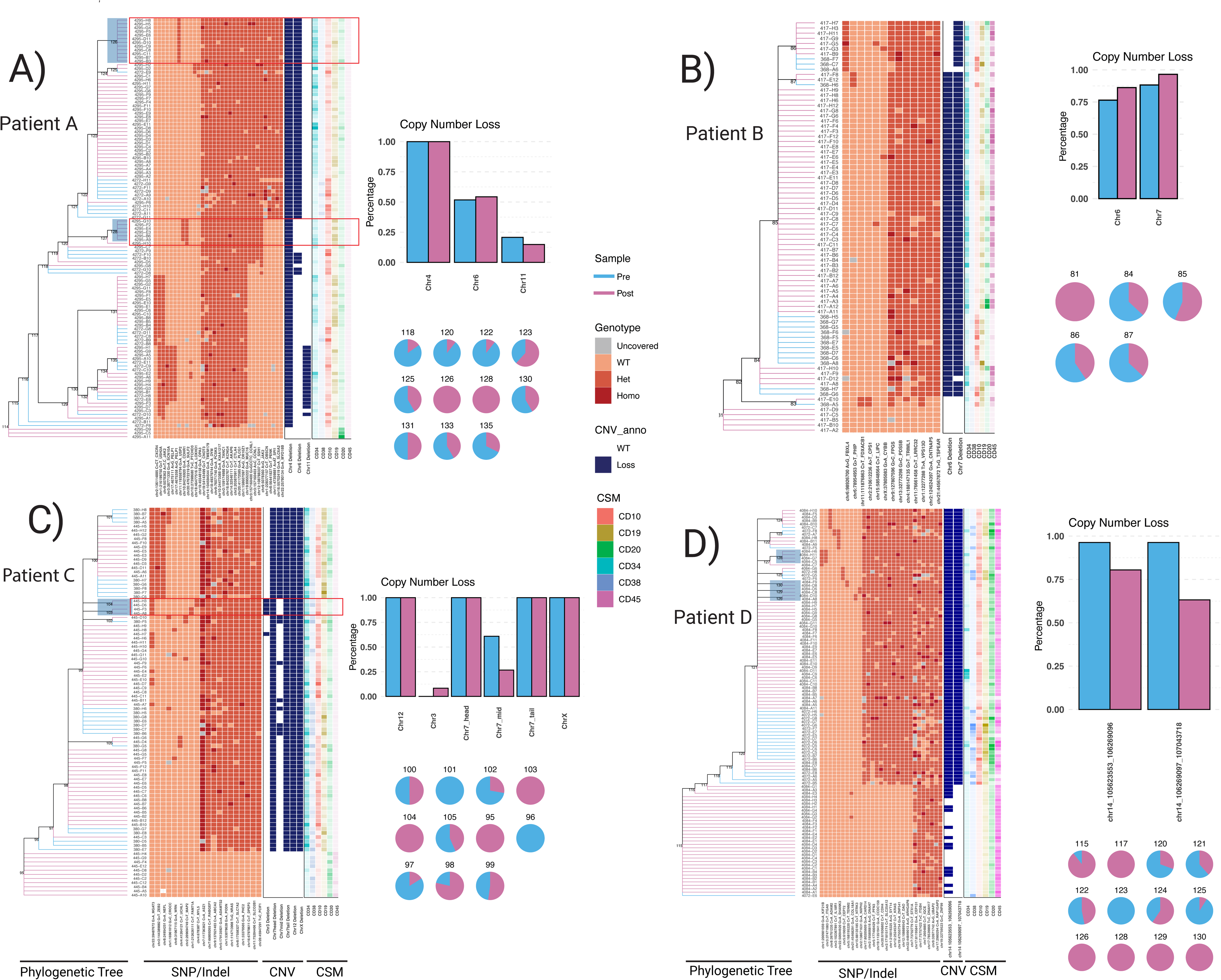
Single-cell sequencing detects clones that undergo selection as patients undergo treatment. Panel A through D depict the phylogenetic trees and associated CNV and surface marker expression. Clones that were not detected in pretreatment are shaded in blue. Relative changes in each recurrent CNV and clonal branch are also depicted for each patient, where it appears that specific SNV are driving the selection of specific clones in patients A and D, while CNV appears to be contributing to a small selected clone in patient C. In contrast to those patients, emergent clones were not identified in patient B.

Post-treatment clones harbored mutations that were not detected in the diagnostic single cells or bulk samples. This includes genes known to be perturbed in treatment-resistant disease^26^, including USP2, which is associated with infant ALL^27^, NTRK3, which occurs in fusions associated with high-risk pALL^28^, and ITGA9, whose expression has been associated with high MRD levels^29^. Other post-treatment associated genes with mutations were CDHR1, FAM71A, KIF21B, MYL5, NXPH2, SULF2, and TENM2. When we examined the emergence of unique CNV in post-treatment cells, patient C had an emergent clone that contained a 90 Mbp chromosome 3q deletion encompassing multiple resistance-association loci. Comparison of our results with 92 relapsed patients revealed these genes mutated in patients that relapse: CDHR1 (4 patients), SULF2 (4), KIF21B (3), NTRK3 (1), ITGA9 (1), and NXPH2 (1) **(Table S3)**^26^.

When we more closely examined the timing of those mutations relative to relapse in that previous study, we also identified patterns of mutation acquisition. ITGA9 and NTRK3 mutations appeared exclusively at relapse, like established relapse-specific resistance mutations in NT5C2 and PRPS1^30^. Conversely, CDHR1, KIF21B, and SULF2 mutations detectable at diagnosis persisted through treatment, resembling genes like CREBBP and NR3C1. The chromosome 3q deletion encompassed focal deletions in BTLA (6/92 relapsed patients) and TBL1XR1/LINC00578 (5/92)^26^. These findings demonstrate that mutations that can drive relapse months to years later are detectable after just four weeks of treatment, providing a window for the early detection of these resistant populations.

## Discussion

Our findings reveal a previously unrecognized role for convergent evolution in generating extensive genetic diversity within pALL patients, changing our understanding of what has been considered a genetically simple cancer. Using PTA-enabled single-cell genome sequencing, we discovered that individual leukemic cells harbor 3,553 mutations, compared to 965 detected in the entire tissue by bulk analysis. This 3.7-fold difference transforms pALL from a genetically simple disease at the tissue level to a much more complex disease at the cellular level. These results also suggest that all cancers are significantly more genetically complex than can be detected with bulk sequencing, and that single-cell sequencing is necessary to accurately measure the true genetic diversity of malignant tissues.

We also found convergent evolution in a significant fraction of patients, suggesting that this underlying genetic complexity can result in multiple clones that evolve to rely on the same pathways for maximal fitness. This includes mutations in KRAS or NRAS in some patients, where we identified up to nine independent ras mutations within a single patient. We also identified convergent evolution of activating JAK2 mutations, as well as clone-specific copy number changes in the same regions in a single patient.

These higher-resolution views of the genetics also enabled us to determine the dynamics of mutational signatures over time. In our index patient, early shared mutations bearing APOBEC signatures (C-to-T and C-to-G at TpC sites) were observed, which contrasts with other patients where APOBEC mutagenesis is a later event. This is consistent with recent studies that have tracked APOBEC signatures in bulk samples from diagnoses through multiple relapses where it was found that APOBEC-associated mutations could rise and fall over time^31^. In contrast, we found a significant association between our de novo signature B that is similar to SBS1 and is associated with deamination of methylated cytosines decreasing in relative frequency over time.

We next sought to connect these complex population genetics with specific phenotypes. By connecting genotype with surface marker expression, we found that chromosome 6 deletions affecting PRDM1, FOXO3, and HDAC2 display increased CD34 and reduced CD20 expression associated with less differentiated B-cells. Similarly, a ZEB2-mutant exhibited altered CD10 expression. By examining primary cells before and after treatment, we were also able to identify clonal populations that increased in frequency after treatment, some of which harbored mutations in genes associated with more aggressive and treatment-resistant disease.

Our approach has several important limitations. First, due to sequencing costs, the number of cells per sample was limited. We do not know how many cells would need to be sequenced to meaningfully saturate our genomic interrogation of clones with specific phenotypes, such as treatment resistance. Similarly, we have only evaluated a limited number of samples. Larger cohorts will be required to identify generalizable genomic patterns of clones that are associated with treatment resistance in pALL patients. Our study establishes the rationale for expanding the number of patients and cells studied with these more sensitive methods. Additionally, our phenotypic annotations were limited to surface marker expression and survival after therapy exposure. PTA has recently been combined with whole transcriptome sequencing from the same cells^32^, which could be evaluated with our recently developed tool PATH, to map entire transcriptomes onto phylogenetic trees to identify changes in the expression of specific genes and pathways that are both heritable and connected to specific phenotypic states ^33^.

This new view of the genetic complexity of pALL has provide new insights into why multi-agent therapy over 2-3 years is essential to cure pALL patients. Our identification of pre-existing clones harboring ITGA9, NTRK3, and SULF2 mutations that survive induction therapy demonstrates that relapse-driving populations can exist at diagnosis below conventional detection thresholds in some patients. This is consistent with the finding that newer higher resolution methods detect cells that are able to survive remission induction therapy in 25-30% of patients. However, these standard MRD methods only detect persistent cells; they do not explain why those cells were able to survive in that patient. Our single-cell approach identifies genes and pathways in those persistent cells that could be used to better risk-stratify and design treatment strategies for patients based on the biology of their persistent disease. In summary, this work begins to reframe our understanding of pALL from a simple oligoclonal disease to a complex, evolutionary ecosystem that we are beginning to deconvolute with high-content single-cell profiling.

## Methods

### Resource Availability

Further information and requests for resources and reagents should be directed to the lead contact, Charles Gawad.

### Data and Code Availability

Single-cell sequencing data have been deposited at SRA at NCBI and are publicly available as of the date of publication. Accession numbers are listed in the key resources table. All original code has been deposited at Mendeley Data and is publicly available as of the date of publication. Any additional information required to reanalyze the data reported in this paper is available from the lead contact upon request.

### Human Subjects

Primary leukemia cells were obtained from pediatric B-acute lymphoblastic leukemia patients aged 1-18 years at St. Jude Children’s Research Hospital (error-corrected sequencing, in vitro treatment on St. Jude Trial TOT16) or Stanford University (scWGA at diagnosis, as well as before/after treatment on COG trial) after IRB approval, as well as informed consent. Samples were collected at diagnosis and after four or six weeks of remission induction therapy. Patient demographics and clinical characteristics are provided in Table S1.

### Cell Lines

DLD-1 colorectal cancer cells (ATCC CCL-221) were maintained in RPMI-1640 (Gibco) supplemented with 10% fetal bovine serum (Gibco) and 1% penicillin/streptomycin (Gibco) at 37°C in 5% CO₂. Medium was changed every 2-3 days, and cells were passaged when reaching 80-90% confluence.

### Sample Collection and Processing

Bone marrow mononuclear cells were isolated using Ficoll-Paque PLUS (GE Healthcare) density gradient centrifugation at 400×g for 30 minutes at 20°C. Cells were resuspended at 1×10⁷ cells/mL in freezing medium containing 90% FBS plus 10% DMSO, frozen in cryovials using controlled-rate freezing, then transferred to liquid nitrogen storage. Cryopreserved samples were thawed using ThawStar system (BioCision) according to manufacturer’s protocol. Cells were washed twice in cold PBS plus 2% FBS, then assessed for viability using trypan blue exclusion. Samples with less than 85% viability were excluded from analysis.

### Single-Cell Isolation and Flow Cytometry

Dead cells were removed using Dead Cell Removal Kit (Miltenyi Biotec). After blocking with 200 μL of 5% rabbit serum (Thermo Fisher Scientific) for 15 minutes at 4°C, cells were stained for 30 minutes at 4°C with the following antibody panel: CD45-APC-H7 (clone 2D1, BD Biosciences) at 5 μL per 10⁶ cells, CD19-APC (clone SJ25C1, BD Biosciences) at 10 μL per 10⁶ cells, CD34-APC-R700 (clone 8G12, BD Biosciences) at 5 μL per 10⁶ cells, CD38-FITC (clone T16, Beckman Coulter) at 10 μL per 10⁶ cells, CD10 (clone HI10A, BD Biosciences) at 5 μL per 10⁶ cells, and CD20-PE (clone L27, BD Biosciences) at 5 μL per 10⁶ cells. Cells were washed twice in PBS plus 2% FBS. MRD blasts were identified as CD45dim/CD19+ cells. Single cells were sorted into 96-well plates containing 3 μL cell buffer (BioSkryb Genomics) using SONY SH800 cell sorter with 130 μm chips. Index sorting parameters recorded cell position, fluorescence intensities, and scatter properties.

### Error-Corrected Sequencing

Adapters with unique molecular identifiers were prepared as previously described (PMID: 25079329). 250 or 500 ng of genomic DNA underwent 30 minutes of chemical fragmentation and library preparation using KAPA HyperPlus Kit (Roche) with adapters containing unique molecular identifiers at 10:1 molar ratio, using 3 μg of adapter per reaction. Unique molecular identifier adapters contained 12-nucleotide random barcodes with forward adapter sequence 5’-ACACTCTTTCCCTACACGACGCTCTTCCGATCT[N12]AGATCGGAAGAGCACACGTCTGAACTCCAGTCA-3’ and reverse adapter sequence 5’-ACACTCTTTCCCTACACGACGCTCTTCCGATCT[N12]AGATCGGAAGAGCGTCGTGTAGGGAAAGAGTGT-3’. PCR amplification and hybrid capture were performed as previously described^14^. Custom capture panel targeted 50 known pediatric acute lymphoblastic leukemia genes with known mutational hotspots based on previous studies^2^. Probe design used IDT xGen Lockdown technology with 2× tiling coverage and 100 bp flanking regions around target sites. Sequencing was performed using MiSeq V2 chemistry with 2×150bp PE reads. Sequences were trimmed to 125 bp using Trimmomatic and unique molecular identifiers placed into headers as previously described using tag_to_header.py^34^. Reads were aligned using BWA ALN with standard parameters. We sorted and indexed using Picard, then performed consensus calling using ConsensusMaker.py with parameters minmem 3, cutoff 0.8, and Ncutoff 0.7. Local realignment was performed using GATK before creating mpileup files. Normal and tumor mpileup files were compared using VarScan Somatic, with somatic mutations requiring P-value less than 10⁻⁴ by Fisher’s exact test^35^. We required germline samples to have fewer than 5 reads with no more than 90% of variant reads on the same DNA strand. Variants underwent annotation with ANNOVAR^36^.

### Single-Cell Exome Sequencing with MDA

Amplified DNA from single cells isolated and whole-genome amplified using Fluidigm C1 System underwent library construction and exome capture with Nextera Rapid Capture Exome Kit (Illumina) according to the manufacturer’s instructions. Exome-enriched libraries underwent sequencing using 2×100 reads on 4 flow cells of HiSeq 2000 or 2500 Sequencing System (Illumina). Adapters were trimmed using Trimmomatic with parameters ILLUMINACLIP:nextera_adapters.fa:2:30:10 TRAILING:25 LEADING:25 SLIDINGWINDOW:4:20 MINLEN:30, followed by alignment with BWA using default parameters. Duplicates were marked using Picard, and local realignment followed by base score recalibration was performed using GATK. Variants were called using GATK and filtered using QD less than 2.0 or FS greater than 60.0 or MQ less than 40.0 or HaplotypeScore greater than 13.0 or MQRankSum less than -12.5 or ReadPosRankSum less than -8.0^37^. On-target coverage was calculated with Picard HsMetrics. Germline SNP locations identified by bulk sequencing were filtered out, followed by the removal of sites identified in normal single cells. For mutation rate estimation, we downsampled the files to 70 million reads, created marked, realigned, and base score-recalibrated BAM files, and then performed variant calling and filtering using the GATK parameters detailed above. Sites found in bulk germline sequencing were subtracted. To account for the background error rate, we subtracted the mean mutation rate in 3 normal cells from each tumor cell.

### Primary Template-Directed Amplification

Single cells underwent whole genome amplification using ResolveDNA WGA Kit v1 (BioSkryb Genomics) per the manufacturer’s instructions. Quality control included assessment by Qubit dsDNA HS assay (ThermoFisher) and Tapestation tracings (Agilent). Libraries were prepared using Illumina DNA Prep Kit with IDT for Illumina DNA/RNA UD Indexes or Kapa HyperPlus workflow (Roche), and some libraries were converted to be compatible with the Ultima Genomics sequencing platform. Size selection targeted 200-500 bp fragments using AMPure XP beads (Beckman Coulter). Libraries were sequenced on NovaSeq 6000 platforms to approximately 8x coverage for whole genome analysis and approximately 60x for exome regions. Pooled libraries for Ultima Genomics were sequenced across two runs, achieving an average 30× depth per cell.

### In Vitro Evolution Validation

An in vitro model of clonal evolution was developed by successively expanding and cloning single DLD-1 (ATCC CCL-221) cells over the course of four months. First, cells from the primary cell culture were expanded in RPMI-1640 with 10% fetal bovine serum for a total of 30 days. A portion of the resulting parental population was cryopreserved (population A), and the remaining cells were used to establish the first filial generation by single-cell cloning. Single-cell cloning was performed by diluting the cells in media at a concentration of <1 cell per 100uL and dispensing 100uL of the suspension into cloning cylinders. Successful clones were expanded over the course of 30 days, periodically increasing the size of the culture dish as the population grew. A portion of expanded clones (N=12) was cryopreserved and another portion was used to establish a second filial generation of cell clones. This procedure of expansion (duration 30-33 days), cryopreservation, and cloning was repeated for a total of 3 filial generations (total time 90 days). The cryopreserved population A and selected clones (N=7) from the first (B,E), second (C) and third (D,F,G,H) filial generations derived from two single cells, were thawed and independently flow sorted into 96-well plates. The isolated cells were subjected to single-cell whole-genome amplification using PTA. PTA products were fragmented and libraries were constructed from the PTA amplification product using Kapa HyperPlus workflow. Libraries resulting from two single-cell amplifications of the parental population (A1 and A2) and two single cells from each of the seven filial clones (B1, B2, E1, E2, C1, C2, D1, D2, G1, G2, F1, F2, H1 and H2) were converted to be compatible with the Ultima Genomics sequencing platform and pooled together on the company facilities on two different runs to reach .an average sequencing depth per cell of 30x. Sequencing data from a single-cell colony derived from the same DLD-1 cell in an independent study was downloaded to use as germline control^20^.

### Variant Calling and Analysis

Somatic variants were called using SCAN2 v1.0 optimized for PTA data^38^, achieving approximately 60-fold fewer false positives per megabase than conventional GATK calling. SCAN2 was run independently for each patient with GRCh38 human genome assembly as reference, Eagle2 panel created with scan2_download_eagle_refpanel.sh for phasing, and dbSNP version 138 for GRCh38 from Google Cloud Storage. The PTA Analysis Toolbox (PTATO) was applied for comparative analyses ^39^. Somatic variants were also called using Sentieon TNscope with matched normal samples. Filtering criteria included quality filters with MQ greater than 59.8, FS less than 5, SOR less than 3, QD 8-20, and ReadPosRankSum greater than or equal to -2. Coverage requirements included a minimum 4× depth and presence in 2 or more cells. Variants in repetitive regions or known artifact sites were excluded. High-quality somatic variants were used for phylogenetic reconstruction with ConDoR using parameters a 0.001, b 0.001, and k 5. Force-calling of low-frequency variants used Mutect2 with force-active true parameter. Mutational burden calculations used mutburden.R script with default parameters. Quality filters removed variants with fewer than 3 supporting reads or less than 5% VAF.

### Mutational Signature Analysis

Mutational signatures were analyzed using SigProfiler suite to create three de novo signatures. SigProfilerMatrixGenerator v1.2 created SBS96 matrices from VCF files. SigProfilerAssignment v0.1 fitted COSMIC signatures v3.4, excluding artifact signatures (27, 43, 45-60, 95). SigProfilerExtractor v1.3 performed de novo extraction with maximum signatures set to 5^18^. Custom scripts extracted sequence context using trinucleotide windows around each variant. Temporal dynamics were assessed by comparing signatures in shared (early) versus unique (late) mutations across cells.

### Copy Number Variation Analysis

CNV detection employed multi-step approach. Shapeit v2 phased heterozygous germline SNPs. Chisel calculated A/B allele read counts. B-allele frequency was computed from count data. Regions with BAF deviation greater than 0.7 or less than 0.3 were classified as LOH. Analysis used 150kb sliding windows with custom haplotype switch minimization. Detection accuracy was validated at 5kb resolution (AUC=0.97). Allelic dropout rate was calculated as percentage of heterozygous germline sites showing only one allele while covered by at least 15 reads.

### Phylogenetic Reconstruction

Phylogenetic trees were constructed using maximum likelihood methods based on somatic SNVs detected across cells. Bootstrap values were calculated with 100 replicates. Branch lengths were correlated with culture time for validation. Linked-read analysis was performed to estimate precision and recall of variant calling as previously described^40^. Phylogenies were time-calibrated using rtreefit with molecular clock assumption. Branch lengths represented mutational distance, converted to time using estimated mutation rates from literature (1.4×10⁻¹⁰ mutations/bp/division).

### Phylogenetic Statistical Analyses

Statistical analyses were performed using R Studio. Phylogenetic signals were assessed using Blomberg’s K statistic with phylosignal package v1.3.0. Significance testing used 999 permutations with p-values less than 0.05 considered significant after Benjamini-Hochberg correction. Growth rates were calculated using Wright-Fisher models with selection in an Approximate Bayesian Computation framework. Simulation parameters included 10,000 iterations and acceptance threshold 0.01. Selection coefficients were calculated from clonal frequencies. BEAST v2.6.3 performed Bayesian phylodynamics analysis for population dynamics. Coalescent theory was applied for clonal timing estimates. Hierarchical clustering used Ward’s method with Euclidean distance. Wright-Fisher simulations modeled clonal dynamics with parameters including population size of 10⁶ to 10⁹ cells, selection coefficients of 0.001 to 0.1 per division, mutation rate of 1×10⁻⁹ per base per division, and bottleneck effects modeled during treatment phases. Phylogenetic trees were visualized using ggtree v3.0.4 with annotations for bootstrap support values, mutational signatures by branch, copy number variants by clade, and surface marker expression levels. Figures were generated using ggplot2, ComplexHeatmap, and custom R scripts. All error bars represent mean ± standard deviation unless otherwise specified.

### Primary Cell Culture and Drug Treatment

Mononuclear cells were isolated using Ficoll-Paque (GE Life Sciences) at sample collection and cryopreserved. One vial per patient was thawed using ThawSTAR system (MedCision) and cultured as previously described^24^. For limited dilution experiments, 750,000 cells were plated per well of 12-well plates and grown for 3 weeks. For drug treatments, 350,000 cells were plated per well of 24-well plates. Medium was changed twice weekly by removing and replacing half volume. Replacement medium included 2× drug concentration for treated samples. All drugs were purchased from Sigma-Aldrich. Concentrations were: mercaptopurine (500, 250, 125, 62.5, and 90 μg/mL), vincristine (810, 162, 32.4, 6.5 μg/mL), daunorubicin (31, 6.2, 1.2, 0.2 μg/mL), prednisolone (concentrations based on published protocols), and asparaginase (19, 9.5, 4.8, 2.4 μg/mL). Live cells were isolated using dead cell removal kit (Miltenyi). DNA was extracted using DNA Universal Kit (Zymo Research), and libraries prepared using HyperPlus Kit (Roche).

#### Temporal Analysis of Treatment Response

Paired samples were collected before treatment and after four weeks of combination chemotherapy (prednisone, vincristine, asparaginase, with daunorubicin for high-risk patients). This timepoint represents when leukemic burden typically decreases from approximately 100 billion to 10-100 million cells (PMID: 32885718). Single-cell analysis was performed on both timepoints to identify treatment-selected clones.

### Data Management and Reproducibility

All analyses were performed using containerized environments for reproducibility with Docker images containing fixed software versions and Conda environments with explicit version specifications. Hardware specifications were documented for compute-intensive analyses. Analysis pipelines were implemented in Snakemake v6.10.0 for workflow management. Custom scripts were written in R v4.1.0 and Python v3.8.0. All code will be deposited in our Github repositories upon publication with detailed documentation and example datasets. Raw sequencing data will be deposited in NCBI SRA with controlled access due to patient privacy requirements. Processed variant matrices and phylogenetic trees will be available through appropriate repositories following journal data sharing policies.

## Acknowledgements

C.G. is supported by the Burroughs Wellcome Fund, Leukemia and Lymphoma Society, Hyundai Hope on Wheels, the American Society of Hematology, NIH Director’s New Innovator Award, and the Chan Zuckerberg Biohub Investigator Program. DAL is supported by the Burroughs Wellcome Fund Career Award for Medical Scientists, the Vallee Scholar Award, the Blood Cancer United Scholar Award and the Mark Foundation Emerging Leader Award. This work was supported by the National Cancer Institute (R33 CA267219), the National Institutes of Health Common Fund Somatic Mosaicism Across Human Tissues (UG3NS132139) and the National Human Genome Research Institute, Center of Excellence in Genomic Science (RM1HG011014). This work was made possible by the MacMillan Family Foundation and the MacMillan Center for the Study of the Non-Coding Cancer Genome at the New York Genome Center. This work was supported by the National Heart Lung and Blood Institute (R01HL157387-01A1). The DLD-1 cell line was a gift from a laboratory at the WCM Belfer Research Building. We thank members of the Gawad and Landau Labs for discussion and advice.

## Data and code availability

Code used for data processing and analysis will be made available upon publication. The data are available through Mendeley Data and have been deposited in SRA at NCBI.

## Competing interests

TP has received conference travel support from BioSkryb. DAL is on the Scientific Advisory Board of Mission Bio, Veracyte, Ultima Genomics and BioSkryb, and has received prior research funding from 10x Genomics, Ultima Genomics, Oxford Nanopore Technologies and Illumina.

## Author Contributions

C.G., D.A.L, Y.P, T.P, I.D., W.E., and V.G. designed research; Y.P, T.P., V.G., A.A., Y.X., J.E., M.M., D.Y., and C.G. performed research; T.P., Y.P., S.R., J.Z., J.Q., J.L., D.K., and M.M. contributed new reagents/analytic tools; Y.P., T.P., C.G., Y.X., S.K., S.R., J.Z., J.Q., J.L., N.O., D.K., M.M., and I.D, analyzed data; and C.G., D.A.L, T.P., Y.P., and W.E. wrote the paper.

## SUPPLEMENTARY MATERIALS

**Table S1.** List of ALL hotspot mutation locations in the error-corrected sequencing capture panel.

**Table S2.** List of recurrent mutations detected in patient SJETV077.

Table S3. List of common mutations detected in relapsed samples in a previous study, highlighting genes that were also selected for in MRD cells as patients underwent treatment.

**Figure S1.** Distribution of duplicate numbers in error-corrected sequencing. Unique molecular identifier family size distributions for (A) germline and (B) leukemia samples.

**Figure S1.** Saturation of sequencing coverage at increasing depth. (A) Exome sequencing of bulk samples reached a saturating coverage breadth of 90% with 10X coverage at 40 million reads. (B) MDA single cells reached a saturating 10X coverage breadth of 55% at 60 million reads.

**Figure S3.** Comparing PTA and MDA exome sequencing at increasing number of reads (A) Fold enrichment (B) Heterozygous SNP sensitivity (C) Coverage at increased depth.

**Figure S4.** Additional ETV6-RUNX1 pALL PTA-based WGS single cell metrics (A) Mutations in bulk and single cells expressed as mutations per Mb (B) Evaluation of genome coverage and variant density for bulk and single cells for additional genomic locations. (D-G) Known SBS signatures that were detected in each sample and single cell, including shared and unique mutations. This is in comparison to the de novo mutation discovery used in Figure 3.

**Figure S5.** Supplementary PTA scWGS Performance Metrics (A) Coverage Median Absolute Deviation (MAD) in neighboring 10Mb-long genomic bins for a different set of samples. Coverage was calculated after downsampling the files at 0.1x. Violin plots are shown when there was more than 1 sample per dataset. Violin plots include mean and standard error bars calculated by computing non-parametric bootstrap. (B) Single-cell coverage breadth at different sequencing efforts for 16 PTA-amplified DLD1 cells. Some library were sequenced twice on a NovaSeq platform. When the same library has been sequenced twice we added the value from the first run and from the combination of the two runs. (C-D). B-allele frequency at germline heterozygous SNPs in chromosome 21 for a PTA-amplified DLD1 cell (C) and for a MDA-amplified NA12878 cell (D). Germline SNPS for the PTA-amplified cell were detected following GATK best practices on a DLD1 single-cell colony. Germline SNPs for the MDA-amplified cell were downloaded from the 1000 Genomes Project. Reads were downsampled in silico and BAF was assessed at any position showing at least 1 read. (E) Allelic dropout (ADO) or percentage of heterozygous germline sites showing only one allele while covered by at least 15 reads. (F) Recall by coverage based on the number of sites in which we obtain the variant allele at heterozygous germline sites. (G) Histogram of the B-allele frequency for the sites showing at least one read along chromosome 21.

**Figure S6.** Additional details, metrics, and validation of single cell variant calling strategy (A) Experimental design and pipeline applied to build phenotypically-annotated PTA phylogenies (B) Precision and recall value ranges provided in external publications and for own data (phycall). (C) Tree representing the possible cell histories under the designed in-vitro evolutionary experiment. Two cells were sampled from the established parental cell population (A1 and A2) whereas the remaining fourteen cells were isolated from seven different single-cell clones from the filial generations. Politomies represent uncertainty in node order given the experimental set up. (D) B-allele frequency of the cells within the patient A phylogeny for the somatic mutations detected in the whole-exome sequencing data. (E) PCA of the fluorescence intensity of the 6 flow-cell markers recorded for the cells within the patient B phylogeny. (F) PCA of the fluorescence intensity of the 6 flow-cell markers recorded for the cells within the patient C phylogeny. (G) Phylodyn analysis for the tumor populations in patients B and C.

## Notes

### Competing Interest Statement

DAL has served as a consultant for Abbvie, AstraZeneca and Illumina, and is on the Scientific Advisory Board of Mission Bio, Pangea, Alethiomics, and C2i Genomics; DAL has received prior research funding from BMS, 10x Genomics, Ultima Genomics, and Illumina unrelated to the current manuscript. CG is a founder and Board Member of BioSkryb Genomics.

### Summary of Updates

The manuscript has been rewritten to focus on the biological and clinical implications of this widespread genetic diversity in pediatric acute lymphoblastic leukemia, rather than the technical advancements that enabled those discoveries.

