## Supplementary figures and images for "Single-Cell Sequencing Reveals Extensive Genetic Diversity Underlying Pediatric ALL Treatment Complexity"

### Figure S1

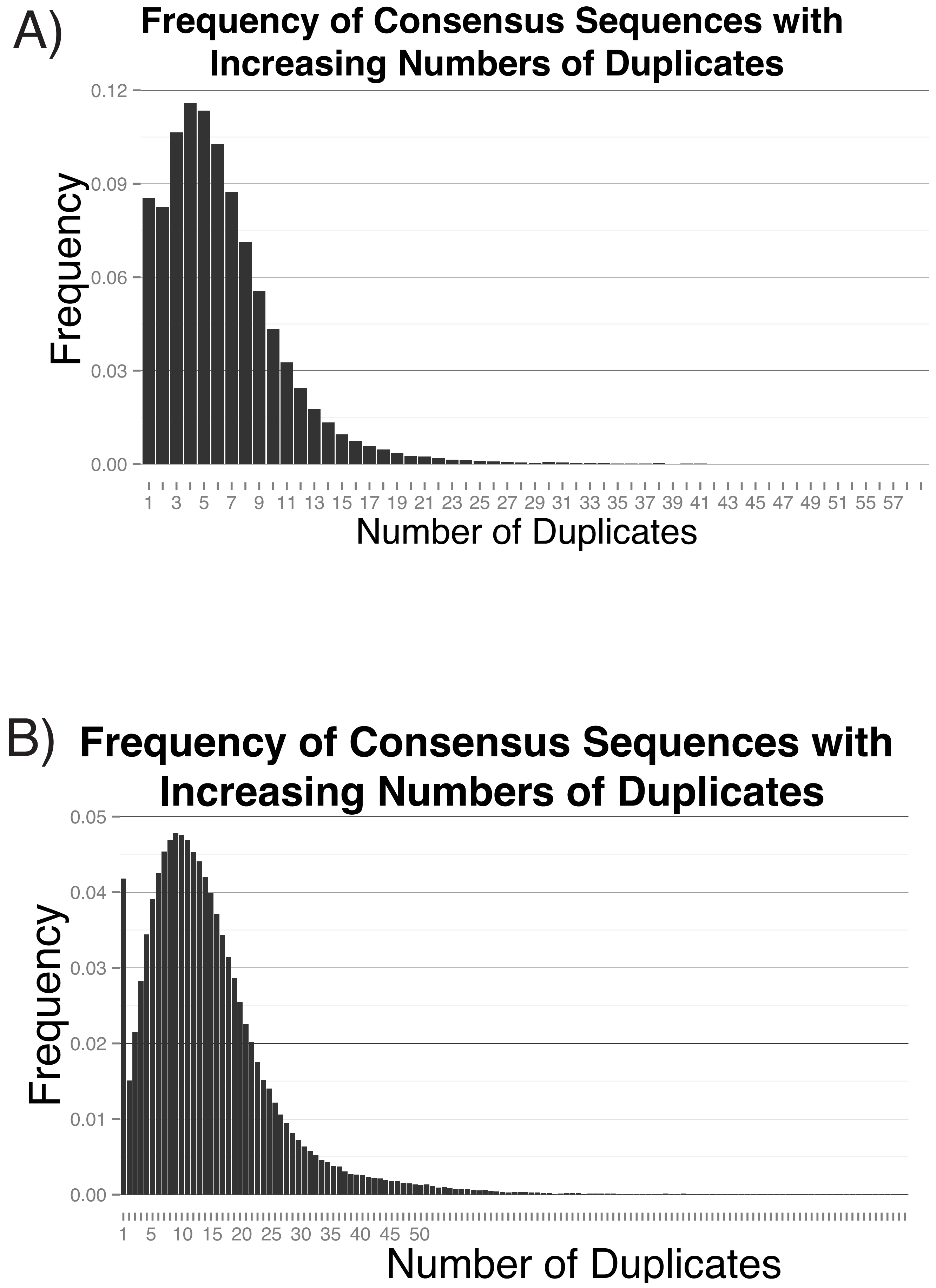

### Figure S2

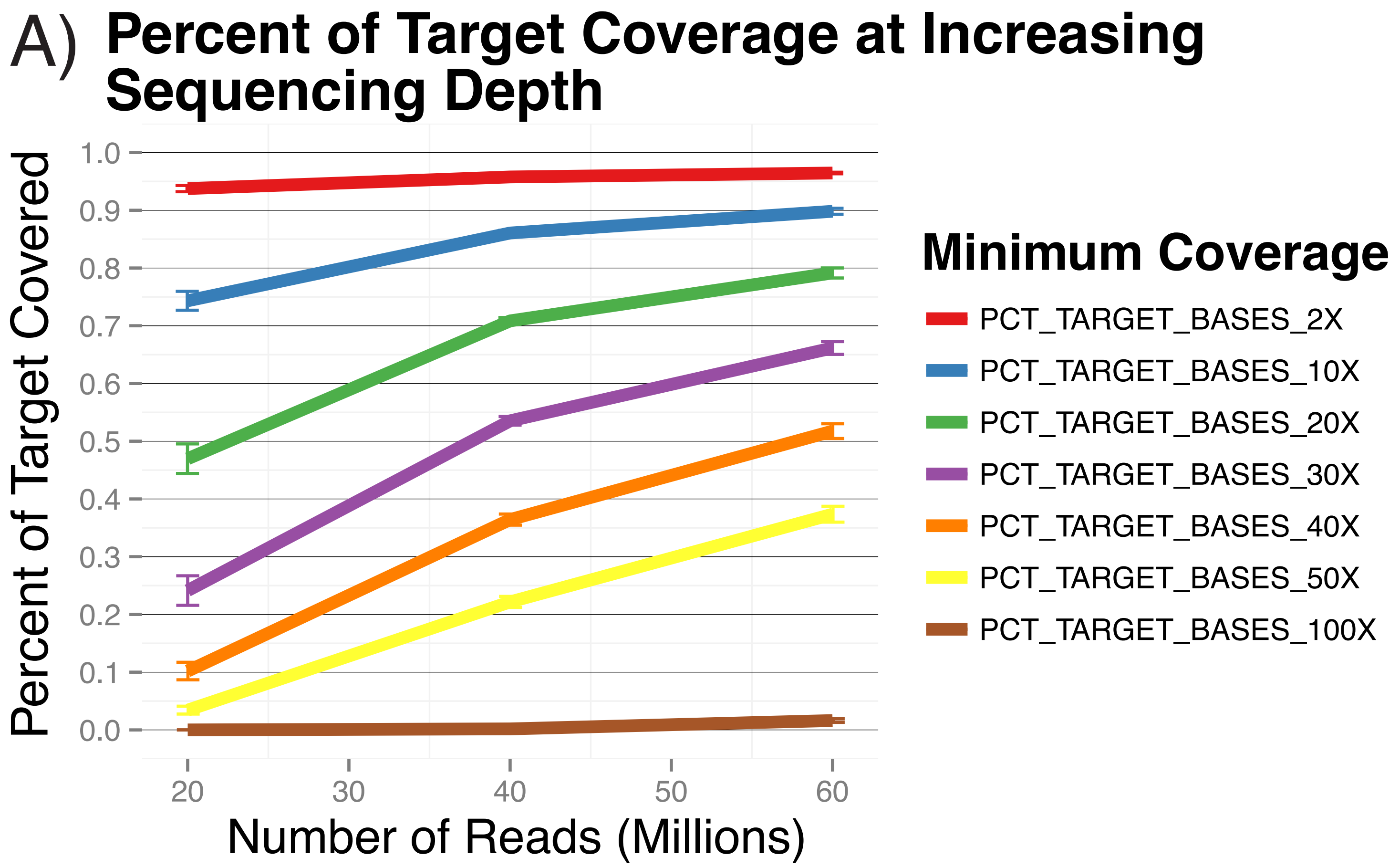

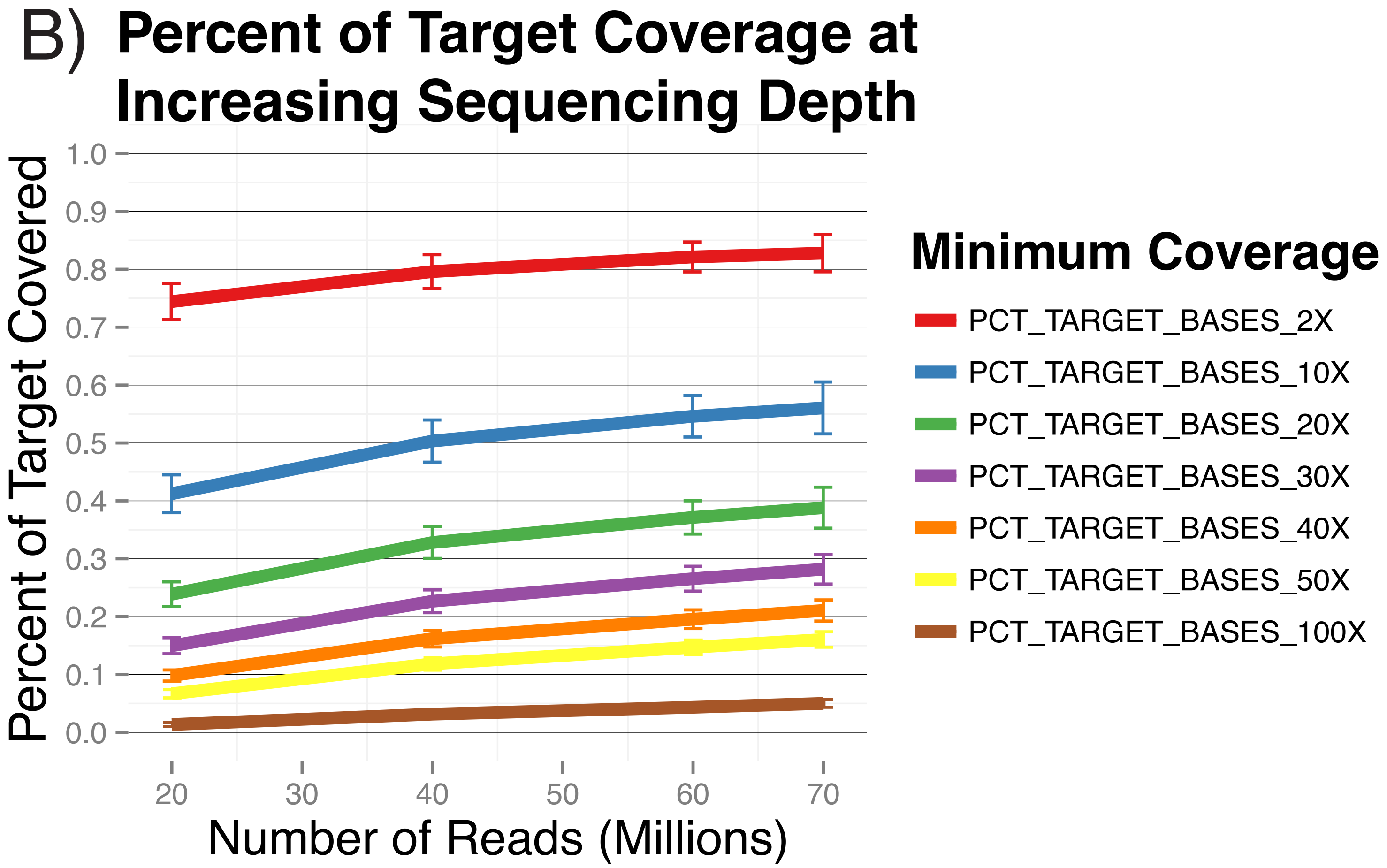

### Figure S3

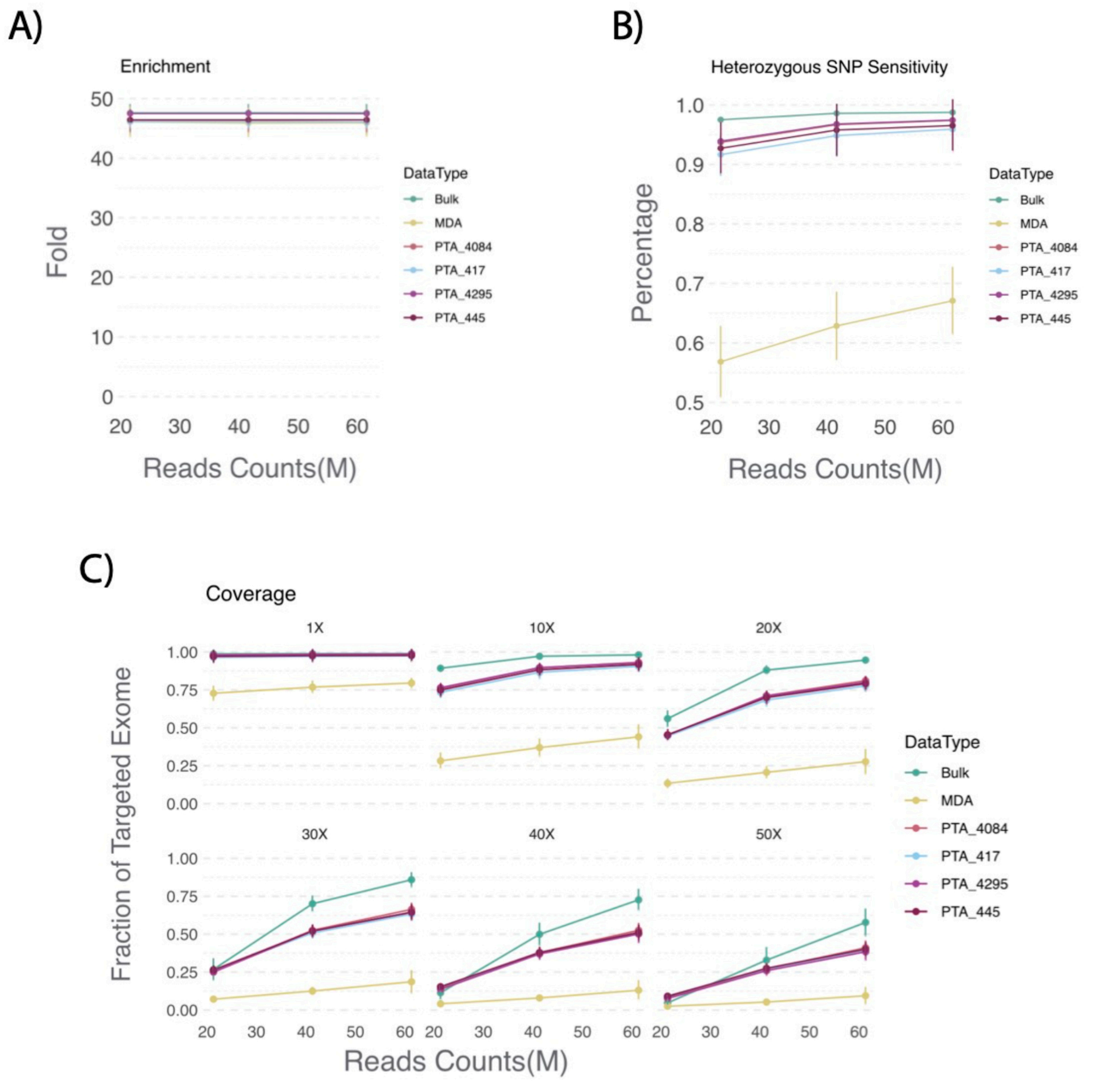

### Figure S4

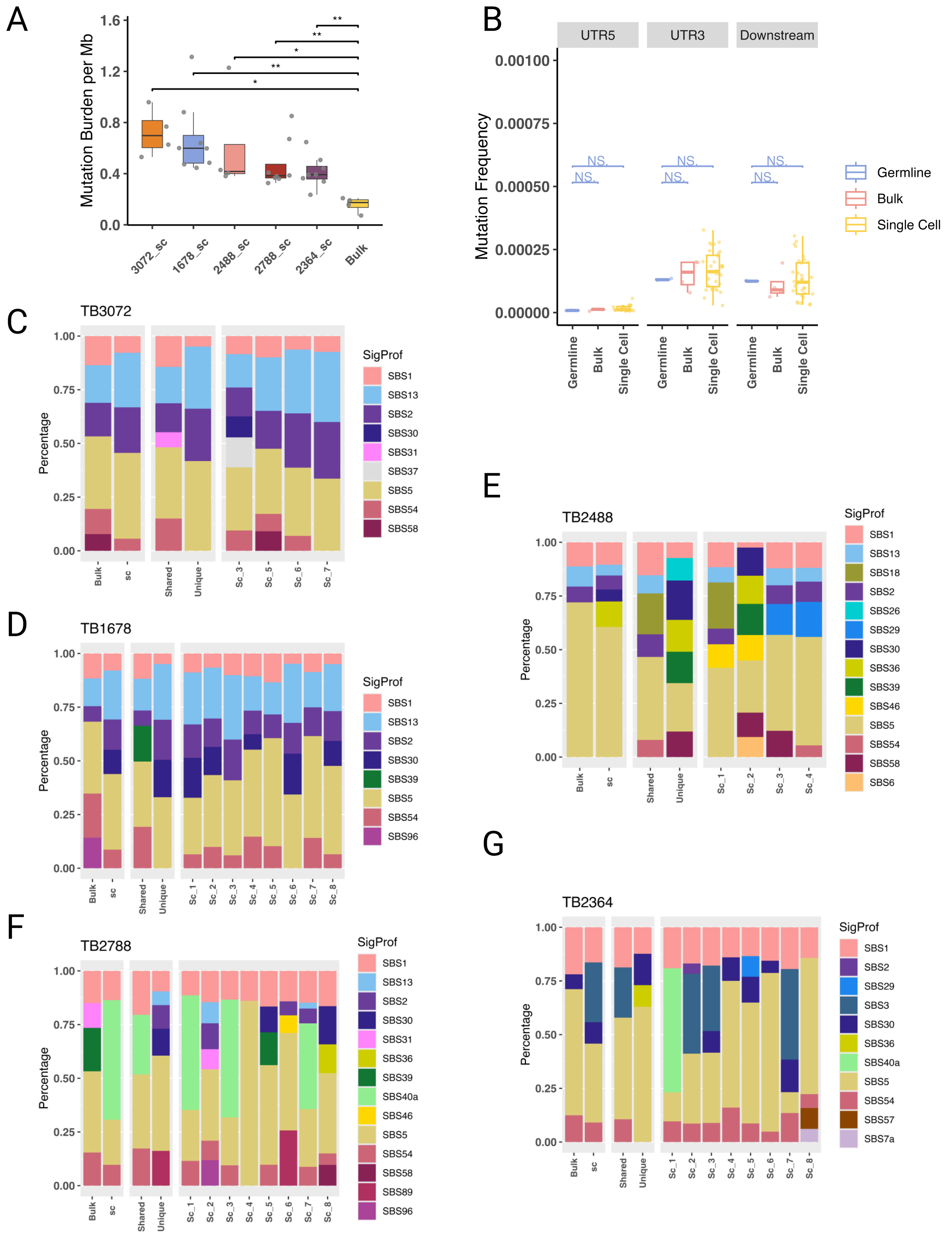

### FIgure S5

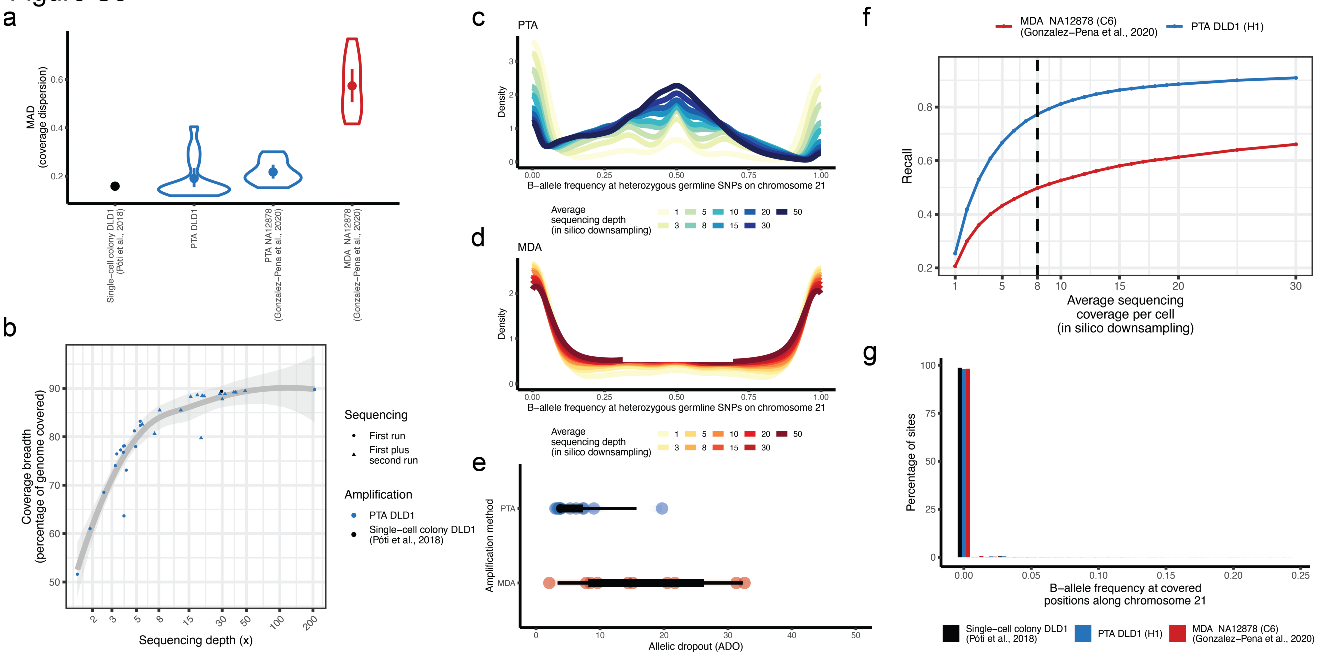

### Figure S6

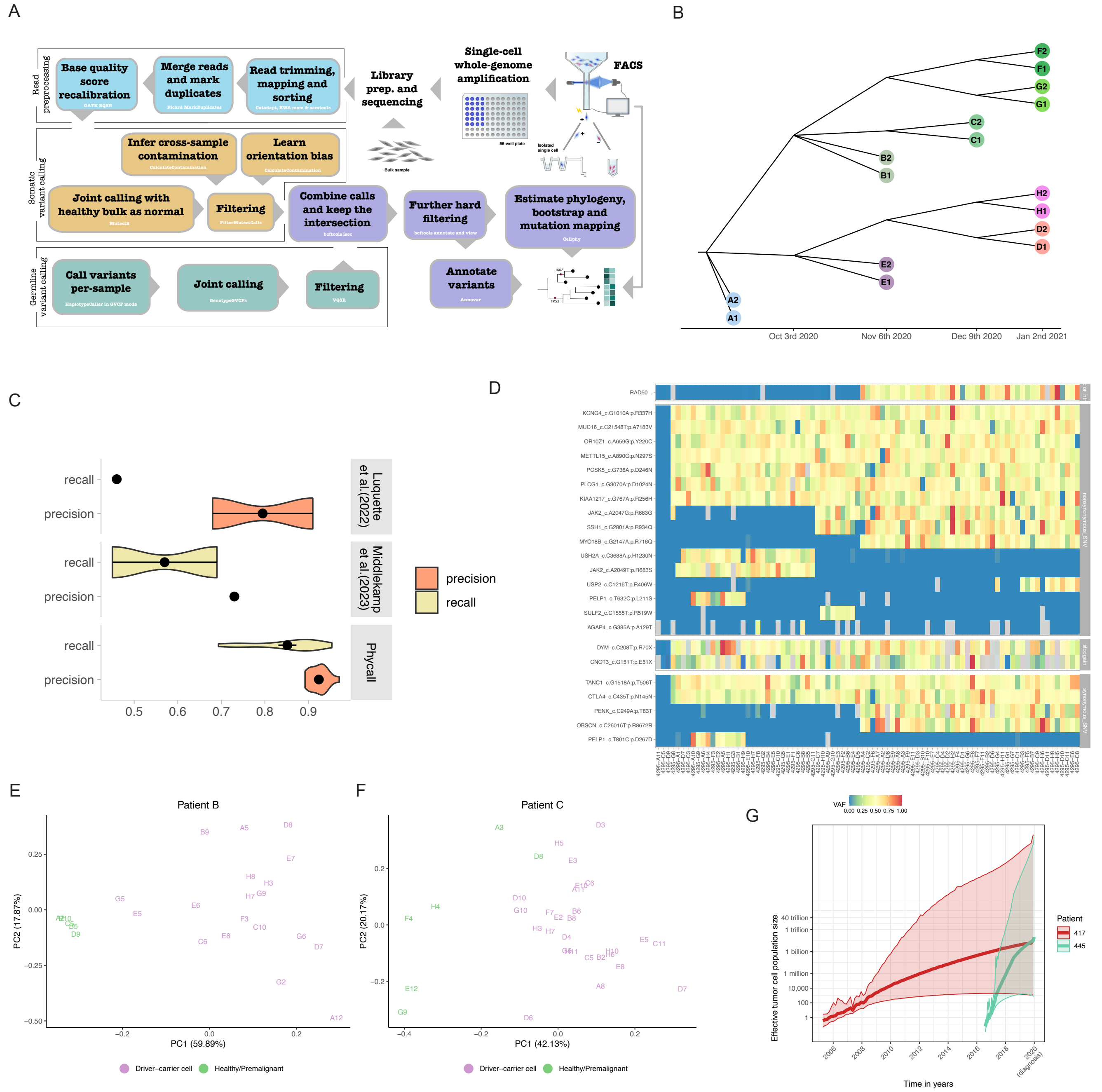
