## Supplementary material for "Single-Cell Sequencing Reveals Extensive Genetic Diversity Underlying Pediatric ALL Treatment Complexity": Table S1

| Chromosome | Start (hg19) | Stop | Gene |
|---|---|---|---|
| 1 | 27099887 | 27100007 | ARID1A |
| 1 | 43814949 | 43815069 | MPL |
| 1 | 65310457 | 65310577 | JAK1 |
| 1 | 115251185 | 115251305 | NRAS |
| 1 | 115252207 | 115252327 | NRAS |
| 1 | 115256470 | 115256590 | NRAS |
| 1 | 115258687 | 115258807 | NRAS |
| 10 | 43600440 | 43600560 | RET |
| 10 | 89692933 | 89693053 | PTEN |
| 10 | 104852901 | 104853021 | NT5C2 |
| 10 | 104857030 | 104857150 | NT5C2 |
| 11 | 533814 | 533934 | HRAS |
| 11 | 32417847 | 32417967 | WT1 |
| 11 | 119148831 | 119148951 | CBL |
| 12 | 25362740 | 25362860 | KRAS |
| 12 | 25368346 | 25368466 | KRAS |
| 12 | 25378548 | 25378668 | KRAS |
| 12 | 25380215 | 25380335 | KRAS |
| 12 | 25398224 | 25398344 | KRAS |
| 12 | 112888150 | 112888270 | PTPN11 |
| 13 | 28592582 | 28592702 | FLT3 |
| 15 | 90631874 | 90631994 | IDH2 |
| 16 | 3788558 | 3788678 | CREBBP |
| 17 | 7577060 | 7577180 | TP53 |
| 17 | 7577478 | 7577598 | TP53 |
| 17 | 7578346 | 7578466 | TP53 |
| 18 | 42531847 | 42531967 | SETBP1 |
| 19 | 17945909 | 17946029 | JAK3 |
| 2 | 25457182 | 25457302 | DNMT3A |
| 2 | 198266774 | 198266894 | SF3B1 |
| 2 | 209113052 | 209113172 | IDH1 |
| 20 | 31022389 | 31022509 | ASXL1 |
| 21 | 36231722 | 36231842 | RUNX1 |
| 3 | 128200670 | 128200790 | GATA2 |
| 4 | 55599261 | 55599381 | KIT |
| 4 | 106156687 | 106156807 | TET2 |
| 4 | 153249324 | 153249444 | FBXW7 |
| 5 | 35873537 | 35873657 | IL7R |
| 5 | 149433585 | 149433705 | CSF1R |
| 5 | 170837487 | 170837607 | NPM1 |
| 7 | 5567936 | 5568056 | ACTB |
| 7 | 103185691 | 103185811 | RELN |
| 7 | 140453076 | 140453196 | BRAF |
| 7 | 148508667 | 148508787 | EZH2 |
| 9 | 5073710 | 5073830 | JAK2 |
| 9 | 133748223 | 133748343 | ABL1 |
| 9 | 139390589 | 139390709 | NOTCH1 |
| 9 | 139399290 | 139399410 | NOTCH1 |

| | | | |
|---|---|---|---|
| X | 1314906 | 1315026 | CRLF2 |
| X | 48649546 | 48649666 | GATA1 |
