## Supplementary material for "Single-Cell Sequencing Reveals Extensive Genetic Diversity Underlying Pediatric ALL Treatment Complexity": Table S2

| Chromosome | Position | Reference Allele | Alternate Allele | Gene Symbol | Change Type | Base and Amino Acid Change |
|---|---|---|---|---|---|---|
| 10 | 100020751 | A | C | LOXL4 | nonsynonymous SNV | LOXL4:NM_032211:exon4:c.T590G:p.M197R |
| 10 | 102783741 | C | T | PDZD7 | nonsynonymous SNV | PDZD7:NM_001195263:exon3:c.G311A:p.G104D,PDZD7:NM_024895:exon3:c.G311A:p.G104D |
| 10 | 25464626 | C | A | GPR158 | nonsynonymous SNV | GPR158:NM_020752:exon1:c.C277A:p.Q93K |
| 10 | 25464650 | G | A | GPR158 | nonsynonymous SNV | GPR158:NM_020752:exon1:c.G301A:p.G101S |
| 10 | 29843823 | C | G | SVIL | nonsynonymous SNV | SVIL:NM_021738:exon5:c.G49C:p.D17H,SVIL:NM_003174:exon7:c.G49C:p.D17H |
| 10 | 30918597 | T | C | LYZL2 | nonsynonymous SNV | LYZL2:NM_183058:exon1:c.A38G:p.K13R |
| 10 | 61833362 | G | A | ANK3 | nonsynonymous SNV | ANK3:NM_020987:exon37:c.C7 |

| | | | | | | |
|---|---|---|---|---|---|---|
| | | | | | | 277T:p.S2426F |
| 10 | 94828146 | T | C | CYP26C1 | nonsynonymous SNV | CYP26C1:NM_183374:exon6:c.T1261C:p.Y421H |
| 10 | 99226266 | C | G | MMS19 | nonsynonymous SNV | MMS19:NM_022362:exon16:c.G1500C:p.Q500H |
| 11 | 118771514 | C | T | BCL9L | nonsynonymous SNV | BCL9L:NM_182557:exon6:c.G2938A:p.G980S |
| 1 | 145299805 | A | C | NBPF10 | nonsynonymous SNV | NBPF10:NM_001039703:exon6:c.A854C:p.E285A |
| 1 | 145299809 | G | A | NBPF10 | nonsynonymous SNV | NBPF10:NM_001039703:exon6:c.G858A:p.M286I |
| 11 | 46880820 | T | A | LRP4 | nonsynonymous SNV | LRP4:NM_002334:exon38:c.A5432T:p.E1811V |
| 11 | 47611491 | T | C | C1QTNF4 | nonsynonymous SNV | C1QTNF4:NM_031909:exon2:c.A872G:p.H291R |
| 1 | 152327810 | C | A | FLG2 | nonsynonymous SNV | FLG2:NM_001014342:exon3:c.G2452T:p.A818S |
| 11 | 55035844 | T | C | TRIM48 | nonsynonymous SNV | TRIM48:NM_024114 |

| | | | | | | |
|---|---|---|---|---|---|---|
| | | | | | | :exon4:c.T574C:p.Y192H |
| 1 | 156347193 | G | A | RHBG | nonsynonymous SNV | RHBG:NM_020407:exon2:c.G289A:p.A97T,RHBG:NM_001256395:exon3:c.G82A:p.A28T,RHBG:NM_001256396:exon3:c.G199A:p.A67T |
| 1 | 156551298 | C | T | TTC24 | stopgain SNV | TTC24:NM_001105669:exon2:c.C142T:p.Q48X |
| 1 | 156622538 | G | A | BCAN | nonsynonymous SNV | BCAN:NM_021948:exon8:c.G1796A:p.R599Q,BCAN:NM_198427:exon8:c.G1796A:p.R599Q |
| 1 | 156843468 | C | G | NTRK1 | nonsynonymous SNV | NTRK1:NM_001012331:exon8:c.C894G:p.H298Q,NTRK1:NM_002529:exon8:c.C894G:p.H298Q,NTRK1:NM_001007792:exon9:c.C804G:p.H268Q |

| | | | | | | |
|---|---|---|---|---|---|---|
| 11 | 57243800 | A | G | RTN4RL2 | nonsynonymous SNV | RTN4RL2:NM_178570:exon3:c.A679G:p.S227G |
| 11 | 57413837 | A | G | YPEL4 | nonsynonymous SNV | YPEL4:NM_145008:exon4:c.T227C:p.L76P |
| 1 | 158641856 | C | G | SPTA1 | nonsynonymous SNV | SPTA1:NM_003126:exon11:c.G1481C:p.R494T |
| 1 | 158641974 | C | G | SPTA1 | nonsynonymous SNV | SPTA1:NM_003126:exon11:c.G1363C:p.D455H |
| 11 | 62296147 | G | C | AHNAK | nonsynonymous SNV | AHNAK:NM_001620:exon5:c.C5742G:p.D1914E |
| 11 | 62296271 | G | A | AHNAK | nonsynonymous SNV | AHNAK:NM_001620:exon5:c.C5618T:p.A1873V |
| 11 | 63671573 | T | A | MARK2 | nonsynonymous SNV | MARK2:NM_001039469:exon15:c.T1630A:p.S544T,MARK2:NM_017490:exon15:c.T1528A:p.S510T |
| 11 | 63884404 | A | G | FLRT1 | nonsynonymous SNV | FLRT1:NM_013280:exon2:c.A665G:p.N222S |

| | | | | | | |
|---|---|---|---|---|---|---|
| 11 | 65321716 | G | C | LTBP3 | nonsynonymous SNV | LTBP3:NM_001130144:exon2:c.C467G:p.P156R,LTBP3:NM_001164266:exon2:c.C116G:p.P39R,LTBP3:NM_021070:exon2:c.C467G:p.P156R |
| 11 | 65325175 | G | T | LTBP3 | nonsynonymous SNV | LTBP3:NM_001130144:exon1:c.C256A:p.Q86K,LTBP3:NM_021070:exon1:c.C256A:p.Q86K |
| 1 | 169572299 | G | A | SELP | nonsynonymous SNV | SELP:NM_003005:exon10:c.C1670T:p.S557L |
| 1 | 186277070 | A | G | PRG4 | nonsynonymous SNV | PRG4:NM_001127710:exon4:c.A1817G:p.E606G,PRG4:NM_001127709:exon5:c.A1940G:p.E647G,PRG4:NM_001127708:exon6:c.A2096G:p.E699G,PRG4:NM_005807 |

| | | | | | | |
|---|---|---|---|---|---|---|
| | | | | | | :exon7:c.A2219G:p.E740G |
| 1 | 186277277 | G | A | PRG4 | nonsynonymous SNV | PRG4:NM_001127710:exon4:c.G2024A:p.G675E,PRG4:NM_001127709:exon5:c.G2147A:p.G716E,PRG4:NM_001127708:exon6:c.G2303A:p.G768E,PRG4:NM_005807:exon7:c.G2426A:p.G809E |
| 1 | 201180415 | A | G | IGFN1 | nonsynonymous SNV | IGFN1:NM_001164586:exon12:c.A6394G:p.T2132A |
| 1 | 201180701 | G | A | IGFN1 | nonsynonymous SNV | IGFN1:NM_001164586:exon12:c.G6680A:p.G2227E |
| 1 | 203821508 | C | G | ZC3H11A | stopgain SNV | ZC3H11A:NM_014827:exon20:c.C2414G:p.S805X |
| 1 | 207787753 | C | T | CR1 | stopgain SNV | CR1:NM_000573:exon32:c.C5230T:p.R1744X,CR1:NM_000651:exon40:c. |

| | | | | | | |
|---|---|---|---|---|---|---|
| | | | | | | C6580T:p.R2194X |
| 1 | 207787796 | T | C | CR1 | nonsynonymous SNV | CR1:NM_000573:exon32:c.T5273C:p.M1758T,CR1:NM_000651:exon40:c.T6623C:p.M2208T |
| 12 | 108961003 | C | T | ISCU | nonsynonymous SNV | ISCU:NM_213595:exon4:c.C377T:p.A126V,ISCU:NM_014301:exon5:c.C302T:p.A101V |
| 12 | 109605831 | C | T | ACACB | nonsynonymous SNV | ACACB:NM_001093:exon3:c.C917T:p.A306V |
| 12 | 112926885 | C | T | PTPN11 | nonsynonymous SNV | PTPN11:NM_002834:exon13:c.C1505T:p.S502L |
| 12 | 11420563 | G | T | PRB3 | nonsynonymous SNV | PRB3:NM_006249:exon3:c.C620A:p.P207Q |
| 12 | 122839007 | C | G | CLIP1 | nonsynonymous SNV | CLIP1:NM_001247997:exon7:c.G1300C:p.E434Q,CLIP1:NM_002956:exon7:c.G1300C:p.E434Q, |

| | | | | | | |
|---|---|---|---|---|---|---|
| | | | | | | CLIP1:NM_198240:exon7:c.G1300C:p.E434Q |
| 12 | 18435367 | C | G | PIK3C2G | nonsynonymous SNV | PIK3C2G:NM_004570:exon2:c.C352G:p.Q118E |
| 12 | 25378562 | C | T | KRAS | nonsynonymous SNV | KRAS:NM_004985:exon4:c.G436A:p.A146T,KRAS:NM_033360:exon4:c.G436A:p.A146T |
| 12 | 25398284 | C | T | KRAS | nonsynonymous SNV | KRAS:NM_004985:exon2:c.G35A:p.G12D,KRAS:NM_033360:exon2:c.G35A:p.G12D |
| 12 | 26276009 | A | C | BHLHE41 | nonsynonymous SNV | BHLHE41:NM_030762:exon5:c.T439G:p.S147A |
| 1 | 226552722 | G | A | PARP1 | nonsynonymous SNV | PARP1:NM_001618:exon19:c.C2639T:p.A880V |
| 12 | 27807681 | G | C | PPFIBP1 | nonsynonymous SNV | PPFIBP1:NM_001198915:exon6:c.G171C:p.L57F,PPFIBP1:NM_001198916: |

| | | | | | | |
|---|---|---|---|---|---|---|
| | | | | | | exon8:c.G630C:p.L210F,PPFIBP1:NM_003622:exon8:c.G630C:p.L210F,PPFIBP1:NM_177444:exon8:c.G630C:p.L210F |
| 1 | 22987700 | G | A | C1QB | nonsynonymous SNV | C1QB:NM_000491:exon3:c.G583A:p.V195I |
| 1 | 22987701 | T | C | C1QB | nonsynonymous SNV | C1QB:NM_000491:exon3:c.T584C:p.V195A |
| 12 | 32138618 | C | G | KIAA1551 | nonsynonymous SNV | KIAA1551:NM_018169:exon4:c.C4729G:p.Q1577E |
| 1 | 24194654 | C | T | FUCA1 | stopgain SNV | FUCA1:NM_000147:exon1:c.G123A:p.W41X |
| 12 | 49434586 | T | C | MLL2 | nonsynonymous SNV | MLL2:NM_003482:exon31:c.A6967G:p.T2323A |
| 12 | 49434613 | T | G | MLL2 | nonsynonymous SNV | MLL2:NM_003482:exon31:c.A6940C:p.T2314P |
| 12 | 50746442 | C | G | FAM186A | nonsynonymous SNV | FAM186A:NM_0011 |

| | | | | | | |
|---|---|---|---|---|---|---|
| | | | | | | 45475:exon4:c.G4173C:p.Q1391H |
| 12 | 50746514 | G | C | FAM186A | nonsynonymous SNV | FAM186A:NM_001145475:exon4:c.C4101G:p.H1367Q |
| 12 | 50747256 | T | C | FAM186A | nonsynonymous SNV | FAM186A:NM_001145475:exon4:c.A3359G:p.E1120G |
| 12 | 57569740 | G | A | LRP1 | nonsynonymous SNV | LRP1:NM_002332:exon24:c.G3842A:p.R1281H |
| 12 | 57648744 | T | C | R3HDM2 | nonsynonymous SNV | R3HDM2:NM_014925:exon22:c.A2743G:p.S915G |
| 12 | 57919829 | C | T | MBD6 | nonsynonymous SNV | MBD6:NM_052897:exon6:c.C1078T:p.R360C |
| 12 | 85413475 | G | A | TSPAN19 | nonsynonymous SNV | TSPAN19:NM_001100917:exon6:c.C380T:p.S127F |
| 12 | 86373909 | G | A | MGAT4C | nonsynonymous SNV | MGAT4C:NM_013244:exon7:c.C595T:p.R199C |
| 13 | 25016769 | G | C | PARP4 | nonsynonymous SNV | PARP4:NM_006437:exon29:c.C3 |

| | | | | | | |
|---|---|---|---|---|---|---|
| | | | | | | 502G:p.L1168V |
| 13 | 28543082 | G | C | CDX2 | nonsynonymous SNV | CDX2:NM_001265:exon1:c.C62G:p.S21C |
| 13 | 33262654 | C | T | PDS5B | nonsynonymous SNV | PDS5B:NM_015032:exon13:c.C1417T:p.R473W |
| 13 | 36049631 | G | C | MAB21L1 | nonsynonymous SNV | MAB21L1:NM_005584:exon1:c.C645G:p.C215W |
| 13 | 37011937 | G | A | CCNA1 | nonsynonymous SNV | CCNA1:NM_001111045:exon3:c.G466A:p.A156T,CCNA1:NM_001111046:exon3:c.G337A:p.A113T,CCNA1:NM_001111047:exon3:c.G337A:p.A113T,CCNA1:NM_003914:exon3:c.G469A:p.A157T |
| 13 | 44455278 | G | A | LACC1 | nonsynonymous SNV | LACC1:NM_001128303:exon2:c.G157A:p.E53K,LACC1:NM_153218:exon2:c.G157A:p.E53K |

| | | | | | | |
|---|---|---|---|---|---|---|
| 13 | 46124044 | C | T | FAM194B | nonsynonymous SNV | FAM194B:NM_182542:exon13:c.G1630A:p.E544K |
| 1 | 36550650 | G | A | TEKT2 | nonsynonymous SNV | TEKT2:NM_014466:exon2:c.G128A:p.R43Q |
| 1 | 38488443 | G | A | UTP11L | nonsynonymous SNV | UTP11L:NM_016037:exon7:c.G640A:p.V214I |
| 14 | 101349407 | A | C | RTL1 | nonsynonymous SNV | RTL1:NM_001134888:exon1:c.T1719G:p.D573E |
| 14 | 105361225 | G | T | CEP170B | nonsynonymous SNV | CEP170B:NM_015005:exon18:c.G4490T:p.R1497L,CEP170B:NM_001112726:exon19:c.G4595T:p.R1532L |
| 14 | 105361230 | A | C | CEP170B | nonsynonymous SNV | CEP170B:NM_015005:exon18:c.A4495C:p.S1499R,CEP170B:NM_001112726:exon19:c.A4600C:p.S1534R |
| 14 | 21899103 | C | T | CHD8 | nonsynonymous SNV | CHD8:NM_001170629:exon1:c |

| | | | | | | |
|---|---|---|---|---|---|---|
| | | | | | | .G700A:p.A234T |
| 14 | 23885450 | G | C | MYH7 | nonsynonymous SNV | MYH7:NM_000257:exon34:c.C4716G:p.I1572M |
| 14 | 23899786 | C | G | MYH7 | nonsynonymous SNV | MYH7:NM_000257:exon11:c.G982C:p.E328Q |
| 14 | 24836309 | T | C | NFATC4 | nonsynonymous SNV | NFATC4:NM_001136022:exon1:c.T49C:p.S17P,NFATC4:NM_001198967:exon1:c.T49C:p.S17P |
| 14 | 32623912 | G | A | ARHGAP5 | nonsynonymous SNV | ARHGAP5:NM_001030055:exon7:c.G4267A:p.E1423K,ARHGAP5:NM_001173:exon7:c.G4264A:p.E1422K |
| 14 | 62462759 | C | T | SYT16 | stopgain SNV | SYT16:NM_031914:exon1:c.C22T:p.Q8X |
| 14 | 91633676 | C | A | C14orf159 | nonsynonymous SNV | C14orf159:NM_001102368:exon4:c.C211A:p.P71T,C14orf159:NM_024952:exon4:c |

| | | | | | | |
|---|---|---|---|---|---|---|
| | | | | | | .C211A:p.P71T,C14orf159:NM_001102367:exon5:c.C211A:p.P71T,C14orf159:NM_001102366:exon6:c.C211A:p.P71T,C14orf159:NM_001102369:exon6:c.C211A:p.P71T |
| 14 | 92472344 | G | C | TRIP11 | stopgain SNV | TRIP11:NM_004239:exon11:c.C1976G:p.S659X |
| 14 | 92922892 | C | T | SLC24A4 | nonsynonymous SNV | SLC24A4:NM_153646:exon12:c.C1195T:p.P399S,SLC24A4:NM_153648:exon12:c.C1003T:p.P335S,SLC24A4:NM_153647:exon13:c.C1138T:p.P380S |
| 15 | 28483865 | G | A | HERC2 | nonsynonymous SNV | HERC2:NM_004667:exon24:c.C3631T:p.R1211C |
| 15 | 43335458 | C | T | UBR1 | nonsynonymous SNV | UBR1:NM_174916:exon15:c.G1 |

| | | | | | | |
|---|---|---|---|---|---|---|
| | | | | | | 804A:p.E602K |
| 15 | 52476793 | G | C | GNB5 | nonsynonymous SNV | GNB5:NM_016194:exon2:c.C81G:p.F27L |
| 15 | 79748677 | C | T | KIAA1024 | nonsynonymous SNV | KIAA1024:NM_015206:exon2:c.C188T:p.T63M |
| 15 | 85188884 | C | G | WDR73 | nonsynonymous SNV | WDR73:NM_032856:exon7:c.G701C:p.G234A |
| 15 | 85383231 | G | A | ALPK3 | nonsynonymous SNV | ALPK3:NM_020778:exon5:c.G1327A:p.E443K |
| 15 | 89400043 | G | C | ACAN | nonsynonymous SNV | ACAN:NM_001135:exon12:c.G4227C:p.E1409D,ACAN:NM_013227:exon12:c.G4227C:p.E1409D |
| 16 | 1265364 | G | A | CACNA1H | nonsynonymous SNV | CACNA1H:NM_001005407:exon28:c.G5144A:p.R1715H,CACNA1H:NM_021098:exon29:c.G5162A:p.R1721H |
| 16 | 1306802 | A | G | TPSD1 | nonsynonymous SNV | TPSD1:NM_012217:e |

| | | | | | | |
|---|---|---|---|---|---|---|
| | | | | | | xon3:c.A259G:p.I87V |
| 16 | 1717967 | C | T | CRAMP1L | nonsynonymous SNV | CRAMP1L:NM_020825:exon17:c.C3107T:p.S1036L |
| 16 | 24834850 | G | A | TNRC6A | nonsynonymous SNV | TNRC6A:NM_014494:exon25:c.G5611A:p.G1871S |
| 16 | 28899031 | G | A | ATP2A1 | nonsynonymous SNV | ATP2A1:NM_004320:exon8:c.G916A:p.A306T,ATP2A1:NM_173201:exon8:c.G916A:p.A306T |
| 16 | 3349644 | G | C | TIGD7 | stopgain SNV | TIGD7:NM_033208:exon2:c.C971G:p.S324X |
| 16 | 3349771 | G | C | TIGD7 | nonsynonymous SNV | TIGD7:NM_033208:exon2:c.C844G:p.L282V |
| 16 | 49670814 | T | A | ZNF423 | nonsynonymous SNV | ZNF423:NM_001271620:exon4:c.A2069T:p.Y690F,ZNF423:NM_015069:exon4:c.A2249T:p.Y750F |
| 16 | 67271614 | C | T | FHOD1 | nonsynonymous SNV | FHOD1:NM_013241 |

| | | | | | | |
|---|---|---|---|---|---|---|
| | | | | | | :exon7:c.G700A:p.E234K |
| 16 | 67663322 | G | C | CTCF | nonsynonymous SNV | CTCF:NM_001191022:exon8:c.G739C:p.D247H,CTCF:NM_006565:exon10:c.G1723C:p.D575H |
| 16 | 72992206 | A | C | ZFHX3 | nonsynonymous SNV | ZFHX3:NM_006885:exon2:c.T1839G:p.D613E |
| 16 | 72992406 | T | C | ZFHX3 | nonsynonymous SNV | ZFHX3:NM_006885:exon2:c.A1639G:p.T547A |
| 16 | 84256031 | G | A | KCNG4 | nonsynonymous SNV | KCNG4:NM_172347:exon3:c.C1352T:p.T451M |
| 16 | 90126823 | T | G | PRDM7 | nonsynonymous SNV | PRDM7:NM_001098173:exon9:c.A1159C:p.M387L |
| 17 | 19607422 | G | A | SLC47A2 | stopgain SNV | SLC47A2:NM_001099646:exon11:c.C979T:p.Q327X,SLC47A2:NM_152908:exon11:c.C1087T:p.Q363X,SLC47A2:NM_00125666 |

| | | | | | | |
|---|---|---|---|---|---|---|
| | | | | | | 3:exon12:c.C1021T:p.Q341X |
| 17 | 21101963 | C | T | TMEM11 | nonsynonymous SNV | TMEM11:NM_003876:exon2:c.G253A:p.A85T |
| 17 | 27028608 | G | A | SUPT6H | nonsynonymous SNV | SUPT6H:NM_003170:exon37:c.G5146A:p.D1716N |
| 17 | 27085444 | C | T | FAM222B | nonsynonymous SNV | FAM222B:NM_001077498:exon3:c.G1533A:p.M511I,FAM222B:NM_018182:exon4:c.G1533A:p.M511I |
| 17 | 27086334 | C | A | FAM222B | nonsynonymous SNV | FAM222B:NM_001077498:exon3:c.G643T:p.A215S,FAM222B:NM_018182:exon4:c.G643T:p.A215S |
| 17 | 3470150 | C | T | TRPV1 | nonsynonymous SNV | TRPV1:NM_080706:exon15:c.G2479A:p.A827T,TRPV1:NM_018727:exon16:c.G2479A:p.A827T,TRPV1:NM_080705:e |

| | | | | | | |
|---|---|---|---|---|---|---|
| | | | | | | xon16:c.G2479A:p.A827T,TRPV1:NM_080704:exon17:c.G2479A:p.A827T |
| 17 | 38249442 | A | G | NR1D1,THRA | nonsynonymous SNV | NR1D1:NM_021724:exon8:c.T1739C:p.L580S,THRA:NM_001190918:exon10:c.A1163G:p.Q388R,THRA:NM_001190919:exon10:c.A1280G:p.Q427R,THRA:NM_003250:exon10:c.A1280G:p.Q427R |
| 17 | 38249525 | G | T | NR1D1,THRA | nonsynonymous SNV | THRA:NM_001190918:exon10:c.G1246T:p.A416S,THRA:NM_001190919:exon10:c.G1363T:p.A455S,THRA:NM_003250:exon10:c.G1363T:p.A455S |
| 17 | 39274422 | C | T | KRTAP4-11 | nonsynonymous SNV | KRTAP4-11:NM_033059:exon |

| | | | | | | |
|---|---|---|---|---|---|---|
| | | | | | | 1:c.G146A:p.C49Y |
| 17 | 41246748 | G | C | BRCA1 | stopgain SNV | BRCA1:NM_007297:exon9:c.C659G:p.S220X,BRCA1:NM_007294:exon10:c.C800G:p.S267X,BRCA1:NM_007300:exon10:c.C800G:p.S267X |
| 17 | 44895945 | C | G | WNT3 | nonsynonymous SNV | WNT3:NM_030753:exon1:c.G19C:p.G7R |
| 17 | 48434506 | C | T | XYLT2 | nonsynonymous SNV | XYLT2:NM_022167:exon9:c.C1834T:p.P612S |
| 17 | 48434579 | G | A | XYLT2 | nonsynonymous SNV | XYLT2:NM_022167:exon9:c.G1907A:p.R636H |
| 1 | 75038329 | G | C | C1orf173 | nonsynonymous SNV | C1orf173:NM_001002912:exon14:c.C3065G:p.A1022G |
| 17 | 5039172 | G | A | USP6 | nonsynonymous SNV | USP6:NM_004505:exon9:c.G613A:p.A205T |
| 17 | 51064116 | G | C | C17orf112 | nonsynonymous SNV | C17orf112:NM_001243552:exo |

| | | | | | | |
|---|---|---|---|---|---|---|
| | | | | | | n3:c.G251C:p.R84T |
| 17 | 7106513 | G | A | DLG4 | nonsynonymous SNV | DLG4:NM_001128827:exon7:c.C632T:p.A211V,DLG4:NM_001365:exon9:c.C770T:p.A257V |
| 17 | 72352973 | C | G | BTBD17 | nonsynonymous SNV | BTBD17:NM_001080466:exon3:c.G1260C:p.Q420H |
| 17 | 72352976 | C | G | BTBD17 | nonsynonymous SNV | BTBD17:NM_001080466:exon3:c.G1257C:p.Q419H |
| 17 | 72353270 | C | G | BTBD17 | nonsynonymous SNV | BTBD17:NM_001080466:exon3:c.G963C:p.W321C |
| 17 | 72353677 | C | G | BTBD17 | nonsynonymous SNV | BTBD17:NM_001080466:exon3:c.G556C:p.E186Q |
| 17 | 72353701 | C | T | BTBD17 | nonsynonymous SNV | BTBD17:NM_001080466:exon3:c.G532A:p.G178S |
| 17 | 72916548 | T | C | USH1G | nonsynonymous SNV | USH1G:NM_173477:exon2:c.A383G:p.N128S |
| 17 | 7577566 | T | C | TP53 | nonsynonymous SNV | TP53:NM_001126115:exon3:c. |

A319G:p.N
107D,TP53
:NM_0011
26116:exo
n3:c.A319
G:p.N107D
,TP53:NM
_0011261
17:exon3:c
.A319G:p.
N107D,TP
53:NM_00
1276697:e
xon3:c.A2
38G:p.N80
D,TP53:N
M_001276
698:exon3
:c.A238G:p
.N80D,TP5
3:NM_001
276699:ex
on3:c.A23
8G:p.N80D
,TP53:NM
_0011261
18:exon6:c
.A598G:p.
N200D,TP
53:NM_00
0546:exon
7:c.A715G:
p.N239D,T
P53:NM_0
01126112:
exon7:c.A
715G:p.N2
39D,TP53:
NM_0011
26113:exo
n7:c.A715
G:p.N239D
,TP53:NM
_0011261

| | | | | | | |
|---|---|---|---|---|---|---|
| | | | | | | 14:exon7:c.A715G:p.N239D,TP53:NM_001276695:exon7:c.A598G:p.N200D,TP53:NM_001276696:exon7:c.A598G:p.N200D,TP53:NM_001276760:exon7:c.A598G:p.N200D,TP53:NM_001276761:exon7:c.A598G:p.N200D |
| 17 | 7579374 | C | A | TP53 | nonsynonymous SNV | TP53:NM_001126118:exon3:c.G196T:p.G66C,TP53:NM_000546:exon4:c.G313T:p.G105C,TP53:NM_001126112:exon4:c.G313T:p.G105C,TP53:NM_001126113:exon4:c.G313T:p.G105C,TP53:NM_001126114:exon4:c.G313T:p.G105C,TP53:N |

| | | | | | | |
|---|---|---|---|---|---|---|
| | | | | | | M_001276695:exon4:c.G196T:p.G66C,TP53:NM_001276696:exon4:c.G196T:p.G66C,TP53:NM_001276760:exon4:c.G196T:p.G66C,TP53:NM_001276761:exon4:c.G196T:p.G66C |
| 1 | 86047232 | G | A | CYR61 | nonsynonymous SNV | CYR61:NM_001554:exon2:c.G248A:p.S83N |
| 18 | 6986227 | G | A | LAMA1 | nonsynonymous SNV | LAMA1:NM_005559:exon37:c.C5288T:p.A1763V |
| 2 | 107049414 | G | A | RGPD3 | nonsynonymous SNV | RGPD3:NM_001144013:exon17:c.C2446T:p.R816C |
| 2 | 109367844 | T | G | RANBP2 | nonsynonymous SNV | RANBP2:NM_006267:exon10:c.T1398G:p.H466Q |
| 2 | 114674544 | A | G | ACTR3 | nonsynonymous SNV | ACTR3:NM_001277140:exon3:c.A31G:p.I11V,ACTR3:NM_005721:exon3:c |

| | | | | | | |
|---|---|---|---|---|---|---|
| | | | | | | .A184G:p.I62V |
| 2 | 132240363 | A | G | TUBA3D | nonsynonymous SNV | TUBA3D:NM_080386:exon5:c.A1295G:p.Y432C |
| 2 | 144381810 | T | C | ARHGAP15 | nonsynonymous SNV | ARHGAP15:NM_018460:exon12:c.T1112C:p.F371S |
| 2 | 144381811 | C | G | ARHGAP15 | nonsynonymous SNV | ARHGAP15:NM_018460:exon12:c.C1113G:p.F371L |
| 2 | 162895664 | C | T | DPP4 | nonsynonymous SNV | DPP4:NM_001935:exon6:c.G397A:p.D133N |
| 2 | 17699517 | G | T | RAD51AP2 | nonsynonymous SNV | RAD51AP2:NM_001099218:exon1:c.C166A:p.R56S |
| 2 | 186668077 | G | A | FSIP2 | nonsynonymous SNV | FSIP2:NM_173651:exon17:c.G14311A:p.E4771K |
| 2 | 198363406 | C | T | HSPD1 | nonsynonymous SNV | HSPD1:NM_002156:exon2:c.G167A:p.G56E,HSPD1:NM_199440:exon2:c.G167A:p.G56E |
| 2 | 207605813 | A | T | MDH1B | nonsynonymous SNV | MDH1B:NM_001039845:exon1 |

| | | | | | | |
|---|---|---|---|---|---|---|
| | | | | | | 0:c.T1424A:p.V475D |
| 2 | 220082421 | C | A | ABCB6 | stopgain SNV | ABCB6:NM_005689:exon2:c.G658T:p.E220X |
| 2 | 220315893 | G | A | SPEG | nonsynonymous SNV | SPEG:NM_005876:exon5:c.G2149A:p.E717K |
| 2 | 26689656 | A | G | OTOF | nonsynonymous SNV | OTOF:NM_194322:exon18:c.T2356C:p.S786P,OTOF:NM_004802:exon19:c.T2125C:p.S709P,OTOF:NM_194323:exon19:c.T2125C:p.S709P,OTOF:NM_194248:exon36:c.T4426C:p.S1476P |
| 2 | 27282122 | A | T | AGBL5 | nonsynonymous SNV | AGBL5:NM_001035507:exon11:c.A1939T:p.S647C,AGBL5:NM_021831:exon11:c.A1939T:p.S647C |
| 2 | 27601820 | C | T | ZNF513 | nonsynonymous SNV | ZNF513:NM_001201459:exon2:c.G127A:p |

| | | | | | | |
|---|---|---|---|---|---|---|
| | | | | | | .A43T,ZNF513:NM_144631:exon3:c.G313A:p.A105T |
| 2 | 73302692 | G | C | RAB11FIP5 | nonsynonymous SNV | RAB11FIP5:NM_015470:exon5:c.C1919G:p.T640S |
| 2 | 75101564 | C | T | HK2 | nonsynonymous SNV | HK2:NM_000189:exon7:c.C863T:p.P288L |
| 2 | 88082426 | C | A | RGPD1,RGPD2 | nonsynonymous SNV | RGPD1:NM_001024457:exon20:c.G4093T:p.D1365Y,RGPD2:NM_001078170:exon20:c.G4117T:p.D1373Y |
| 3 | 102196305 | C | T | ZPLD1 | nonsynonymous SNV | ZPLD1:NM_175056:exon11:c.C1139T:p.P380L |
| 3 | 127783850 | C | G | SEC61A1 | nonsynonymous SNV | SEC61A1:NM_013336:exon8:c.C747G:p.I249M |
| 3 | 130799347 | C | T | NEK11 | nonsynonymous SNV | NEK11:NM_001146003:exon3:c.C251T:p.S84F,NEK11:NM_024800:exon4:c.C251T:p.S84F,NEK11 |

| | | | | | | |
|---|---|---|---|---|---|---|
| | | | | | | :NM_145910:exon4:c.C251T:p.S84F |
| 3 | 168833952 | A | G | MECOM | nonsynonymous SNV | MECOM:NM_001105078:exon7:c.T1144C:p.F382L,MECOM:NM_001163999:exon7:c.T1147C:p.F383L,MECOM:NM_001164000:exon7:c.T1144C:p.F382L,MECOM:NM_001205194:exon7:c.T1144C:p.F382L,MECOM:NM_005241:exon7:c.T1144C:p.F382L,MECOM:NM_001105077:exon8:c.T1339C:p.F447L,MECOM:NM_004991:exon8:c.T1708C:p.F570L |
| 3 | 168834012 | A | G | MECOM | nonsynonymous SNV | MECOM:NM_001105078:exon7:c.T1084C:p.S362P,MECOM:NM |

| | | | | | | |
|---|---|---|---|---|---|---|
| | | | | | | _001163999:exon7:c.T1087C:p.S363P,MECOM:NM_001164000:exon7:c.T1084C:p.S362P,MECOM:NM_001205194:exon7:c.T1084C:p.S362P,MECOM:NM_005241:exon7:c.T1084C:p.S362P,MECOM:NM_001105077:exon8:c.T1279C:p.S427P,MECOM:NM_004991:exon8:c.T1648C:p.S550P |
| 3 | 183368618 | C | G | KLHL24 | nonsynonymous SNV | KLHL24:NM_017644:exon3:c.C474G:p.F158L |
| 3 | 183689545 | A | C | ABCC5 | nonsynonymous SNV | ABCC5:NM_005688:exon11:c.T1567G:p.S523A |
| 3 | 32031962 | A | G | ZNF860 | nonsynonymous SNV | ZNF860:NM_001137674:exon2:c.A1391G:p.H464R |

| | | | | | | |
|---|---|---|---|---|---|---|
| 3 | 32032046 | G | A | ZNF860 | nonsynonymous SNV | ZNF860:NM_001137674:exon2:c.G1475A:p.R492H |
| 3 | 33155909 | C | T | CRTAP | nonsynonymous SNV | CRTAP:NM_006371:exon1:c.C340T:p.R114C |
| 3 | 57443708 | C | T | DNAH12 | nonsynonymous SNV | DNAH12:NM_178504:exon22:c.G3007A:p.E1003K |
| 4 | 100234991 | C | T | ADH1B | nonsynonymous SNV | ADH1B:NM_000668:exon6:c.G815A:p.R272Q |
| 4 | 146062683 | A | G | OTUD4 | nonsynonymous SNV | OTUD4:NM_001102653:exon19:c.T1736C:p.I579T |
| 4 | 15004680 | T | A | CPEB2 | nonsynonymous SNV | CPEB2:NM_001177381:exon1:c.T383A:p.I128N,CPEB2:NM_001177382:exon1:c.T383A:p.I128N,CPEB2:NM_001177383:exon1:c.T383A:p.I128N,CPEB2:NM_001177384:exon1:c.T383A:p.I128N,CPEB2:NM |

| | | | | | | |
|---|---|---|---|---|---|---|
| | | | | | | _182485:exon1:c.T383A:p.I128N,CPEB2:NM_182646:exon1:c.T383A:p.I128N |
| 4 | 15004758 | C | G | CPEB2 | nonsynonymous SNV | CPEB2:NM_001177381:exon1:c.C461G:p.S154C,CPEB2:NM_001177382:exon1:c.C461G:p.S154C,CPEB2:NM_001177383:exon1:c.C461G:p.S154C,CPEB2:NM_001177384:exon1:c.C461G:p.S154C,CPEB2:NM_182485:exon1:c.C461G:p.S154C,CPEB2:NM_182646:exon1:c.C461G:p.S154C |
| 4 | 1918753 | G | A | WHSC1 | nonsynonymous SNV | WHSC1:NM_133334:exon3:c.G916A:p.E306K,WHSC1:NM_001042424:exon4:c.G916A:p.E306 |

| | | | | | | |
|---|---|---|---|---|---|---|
| | | | | | | K,WHSC1:NM_133335:exon4:c.G916A:p.E306K,WHSC1:NM_133331:exon5:c.G916A:p.E306K,WHSC1:NM_007331:exon6:c.G916A:p.E306K,WHSC1:NM_133330:exon6:c.G916A:p.E306K |
| 4 | 1920348 | G | A | WHSC1 | nonsynonymous SNV | WHSC1:NM_133334:exon4:c.G1408A:p.E470K,WHSC1:NM_001042424:exon5:c.G1408A:p.E470K,WHSC1:NM_133335:exon5:c.G1408A:p.E470K,WHSC1:NM_133331:exon6:c.G1408A:p.E470K,WHSC1:NM_007331:exon7:c.G1408A:p.E470K,WHSC1:NM_133330:exon7:c.G14 |

| | | | | | | |
|---|---|---|---|---|---|---|
| | | | | | | 08A:p.E470K |
| 4 | 3254966 | C | T | MSANTD1 | nonsynonymous SNV | MSANTD1:NM_001042690:exon2:c.C353T:p.A118V |
| 4 | 3255001 | G | A | MSANTD1 | nonsynonymous SNV | MSANTD1:NM_001042690:exon2:c.G388A:p.G130R |
| 4 | 99355110 | G | C | RAP1GDS1 | nonsynonymous SNV | RAP1GDS1:NM_001100430:exon11:c.G1191C:p.Q397H,RAP1GDS1:NM_001100428:exon12:c.G1320C:p.Q440H,RAP1GDS1:NM_001100429:exon12:c.G1317C:p.Q439H,RAP1GDS1:NM_001100426:exon13:c.G1467C:p.Q489H,RAP1GDS1:NM_001100427:exon13:c.G1464C:p.Q488H,RAP1GDS1:NM_021159:exon13:c.G1464C:p.Q488H |

| | | | | | | |
|---|---|---|---|---|---|---|
| 5 | 112151253 | C | G | APC | nonsynonymous SNV | APC:NM_001127511:exon7:c.C842G:p.S281C,APC:NM_000038:exon9:c.C896G:p.S299C,APC:NM_001127510:exon10:c.C896G:p.S299C |
| 5 | 140250188 | C | G | PCDHA11 | nonsynonymous SNV | PCDHA11:NM_018902:exon1:c.C1500G:p.D500E,PCDHA11:NM_031861:exon1:c.C1500G:p.D500E |
| 5 | 140517305 | C | G | PCDHB5 | nonsynonymous SNV | PCDHB5:NM_015669:exon1:c.C2289G:p.F763L |
| 5 | 140812343 | G | A | PCDHGA12 | nonsynonymous SNV | PCDHGA12:NM_003735:exon1:c.G2017A:p.A673T,PCDHGA12:NM_032094:exon1:c.G2017A:p.A673T |
| 5 | 157170792 | G | C | LSM11 | nonsynonymous SNV | LSM11:NM_173491:exon1:c.G34C:p.G12R |
| 5 | 159545858 | A | G | PWWP2A | nonsynonymous SNV | PWWP2A:NM_0011 |

| | | | | | | |
|---|---|---|---|---|---|---|
| | | | | | | 30864:exon1:c.T538C:p.F180L,PWWP2A:NM_001267035:exon1:c.T538C:p.F180L,PWWP2A:NM_052927:exon1:c.T538C:p.F180L |
| 5 | 228322 | A | G | SDHA | nonsynonymous SNV | SDHA:NM_004168:exon6:c.A644G:p.Y215C |
| 5 | 23527167 | G | C | PRDM9 | nonsynonymous SNV | PRDM9:NM_020227:exon11:c.G1970C:p.R657T |
| 5 | 23527472 | G | A | PRDM9 | nonsynonymous SNV | PRDM9:NM_020227:exon11:c.G2275A:p.D759N |
| 5 | 23527637 | A | C | PRDM9 | nonsynonymous SNV | PRDM9:NM_020227:exon11:c.A2440C:p.S814R |
| 5 | 23527724 | A | G | PRDM9 | nonsynonymous SNV | PRDM9:NM_020227:exon11:c.A2527G:p.N843D |
| 5 | 32089885 | G | A | PDZD2 | nonsynonymous SNV | PDZD2:NM_178140:exon19:c.G6331A:p.E2111K |
| 5 | 40692166 | C | A | PTGER4 | nonsynonymous SNV | PTGER4:NM_000958 |

| | | | | | | |
|---|---|---|---|---|---|---|
| | | | | | | :exon3:c.C1153A:p.L385M |
| 5 | 52365978 | C | T | ITGA2 | nonsynonymous SNV | ITGA2:NM_002203:exon17:c.C2123T:p.S708L |
| 5 | 79616400 | G | A | SPZ1 | nonsynonymous SNV | SPZ1:NM_032567:exon1:c.G366A:p.M122I |
| 6 | 107955780 | A | G | SOBP | nonsynonymous SNV | SOBP:NM_018013:exon6:c.A1732G:p.S578G |
| 6 | 12126032 | C | G | HIVEP1 | nonsynonymous SNV | HIVEP1:NM_002114:exon4:c.C6004G:p.Q2002E |
| 6 | 1390534 | A | G | FOXF2 | nonsynonymous SNV | FOXF2:NM_001452:exon1:c.A352G:p.S118G |
| 6 | 1390535 | G | C | FOXF2 | nonsynonymous SNV | FOXF2:NM_001452:exon1:c.G353C:p.S118T |
| 6 | 168377119 | T | C | HGC6.3 | nonsynonymous SNV | HGC6.3:NM_001129895:exon1:c.A214G:p.M72V |
| 6 | 168430272 | T | A | KIF25 | nonsynonymous SNV | KIF25:NM_005355:exon2:c.T7A:p.W3R,KIF25:NM_030615:exon |

| | | | | | | |
|---|---|---|---|---|---|---|
| | | | | | | 2:c.T7A:p.W3R |
| 6 | 31605302 | G | A | PRRC2A | nonsynonymous SNV | PRRC2A:NM_004638:exon31:c.G6413A:p.R2138Q,PRRC2A:NM_080686:exon31:c.G6413A:p.R2138Q |
| 6 | 33415660 | G | A | SYNGAP1 | nonsynonymous SNV | SYNGAP1:NM_006772:exon18:c.G3835A:p.A1279T |
| 6 | 6002600 | G | C | NRN1 | nonsynonymous SNV | NRN1:NM_016588:exon2:c.C186G:p.I62M |
| 6 | 89895130 | T | C | GABRR1 | nonsynonymous SNV | GABRR1:NM_001256703:exon6:c.A644G:p.K215R,GABRR1:NM_002042:exon7:c.A695G:p.K232R,GABRR1:NM_001256704:exon8:c.A434G:p.K145R,GABRR1:NM_001267582:exon9:c.A434G:p.K145R |
| 7 | 100482090 | G | C | SRRT | nonsynonymous SNV | SRRT:NM_001128852:exon7:c.G859C:p.E |

| | | | | | | |
|---|---|---|---|---|---|---|
| | | | | | | 287Q,SRRT:NM_001128853:exon7:c.G859C:p.E287Q,SRRT:NM_001128854:exon7:c.G859C:p.E287Q,SRRT:NM_015908:exon7:c.G859C:p.E287Q |
| 7 | 105254947 | G | C | ATXN7L1 | nonsynonymous SNV | ATXN7L1:NM_138495:exon8:c.C1462G:p.P488A,ATXN7L1:NM_020725:exon10:c.C1834G:p.P612A |
| 7 | 105254977 | A | G | ATXN7L1 | nonsynonymous SNV | ATXN7L1:NM_138495:exon8:c.T1432C:p.S478P,ATXN7L1:NM_020725:exon10:c.T1804C:p.S602P |
| 7 | 150094143 | T | C | ZNF775 | nonsynonymous SNV | ZNF775:NM_173680:exon3:c.T574C:p.S192P |
| 7 | 150768907 | G | C | SLC4A2 | nonsynonymous SNV | SLC4A2:NM_001199693:exon14:c.G2296C:p.E766Q, |

| | | | | | | |
|---|---|---|---|---|---|---|
| | | | | | | SLC4A2:NM_001199694:exon14:c.G2281C:p.E761Q, SLC4A2:NM_001199692:exon15:c.G2323C:p.E775Q, SLC4A2:NM_003040:exon15:c.G2323C:p.E775Q |
| 7 | 151962134 | G | T | MLL3 | stopgain SNV | MLL3:NM_170606:exon8:c.C1173A:p.C391X |
| 7 | 2581772 | G | C | BRAT1 | nonsynonymous SNV | BRAT1:NM_152743:exon7:c.C997G:p.Q333E |
| 7 | 27238961 | T | C | HOXA13 | nonsynonymous SNV | HOXA13:NM_000522:exon1:c.A736G:p.M246V |
| 7 | 27238988 | C | G | HOXA13 | nonsynonymous SNV | HOXA13:NM_000522:exon1:c.G709C:p.A237P |
| 7 | 33044951 | C | G | FKBP9 | nonsynonymous SNV | FKBP9:NM_007270:exon10:c.C1701G:p.H567Q |
| 7 | 33044951 | C | T | FKBP9 | synonymous SNV | FKBP9:NM_007270:exon10:c.C1 |

| | | | | | | |
|---|---|---|---|---|---|---|
| | | | | | | 701T:p.H567H |
| 7 | 57187725 | G | T | ZNF479 | nonsynonymous SNV | ZNF479:NM_033273:exon5:c.C1397A:p.T466K |
| 7 | 57187780 | A | C | ZNF479 | nonsynonymous SNV | ZNF479:NM_033273:exon5:c.T1342G:p.L448V |
| 7 | 75613144 | G | A | POR | nonsynonymous SNV | POR:NM_000941:exon10:c.G1036A:p.D346N |
| 7 | 89790570 | G | C | STEAP1 | nonsynonymous SNV | STEAP1:NM_012449:exon3:c.G536C:p.S179T |
| 8 | 143695347 | A | T | ARC | nonsynonymous SNV | ARC:NM_015193:exon1:c.T286A:p.C96S |
| 8 | 145059786 | T | A | PARP10 | nonsynonymous SNV | PARP10:NM_032789:exon4:c.A467T:p.N156I |
| 8 | 26372007 | T | A | DPYSL2 | nonsynonymous SNV | DPYSL2:NM_001197293:exon1:c.T166A:p.S56T |
| 9 | 113341430 | A | T | SVEP1 | nonsynonymous SNV | SVEP1:NM_153366:exon1:c.T394A:p.Y132N |
| 9 | 12702433 | T | C | TYRP1 | nonsynonymous SNV | TYRP1:NM_000550:exon5:c.T10 |

| | | | | | | |
|---|---|---|---|---|---|---|
| | | | | | | 76C:p.V359A |
| 9 | 133944381 | A | T | LAMC3 | nonsynonymous SNV | LAMC3:NM_006059:exon16:c.A2834T:p.Q945L |
| 9 | 135139870 | A | T | SETX | nonsynonymous SNV | SETX:NM_015046:exon26:c.T7790A:p.L2597Q |
| 9 | 135946015 | T | C | CEL | nonsynonymous SNV | CEL:NM_001807:exon10:c.T1463C:p.I488T |
| 9 | 139740877 | C | G | C9orf172 | nonsynonymous SNV | C9orf172:NM_001080482:exon1:c.C2011G:p.R671G |
| 9 | 139992338 | G | A | MAN1B1 | nonsynonymous SNV | MAN1B1:NM_016219:exon5:c.G679A:p.E227K |
| 9 | 140080774 | C | T | ANAPC2 | nonsynonymous SNV | ANAPC2:NM_013366:exon3:c.G775A:p.E259K |
| 9 | 34089478 | C | G | DCAF12 | nonsynonymous SNV | DCAF12:NM_015397:exon8:c.G1135C:p.A379P |
| 9 | 34990713 | C | T | DNAJB5 | nonsynonymous SNV | DNAJB5:NM_001135005:exon1:c.C86T:p.S29F,DNAJB5:NM_001135004:ex |

on2:c.C21
2T:p.S71F
