## Supplementary material for "Single-Cell Sequencing Reveals Extensive Genetic Diversity Underlying Pediatric ALL Treatment Complexity": Table S3

Rows in blue are genes that were found to harbor MRD-specific variants in our study.

| Number of Samples | Gene | Mutation Type | Change from Diagnosis to Relapse |
|---|---|---|---|
| 1 | CDHR1 | missense | Acquired |
| 1 | CDHR1 | silent | Acquired |
| 1 | CDHR1 | UTR_3 | Acquired |
| 1 | CDHR1 | UTR_3 | Lost |
| 1 | CDHR1 | UTR_3 | Preserved from subclone |
| 1 | CREBBP | frameshift | Acquired |
| 2 | CREBBP | missense | Acquired |
| 2 | CREBBP | nonsense | Acquired |
| 1 | CREBBP | missense | Lost |
| 1 | CREBBP | silent | NA |
| 2 | CREBBP | missense | Preserved |
| 2 | CREBBP | proteinDel | Preserved |
| 3 | CREBBP | missense | Preserved from subclone |
| 1 | CREBBP | nonsense | Preserved from subclone |
| 1 | CREBBP | proteinDel | Preserved from subclone |
| 1 | CREBBP | silent | Preserved from subclone |
| 2 | CREBBP | missense | Preserved gain allele |
| 1 | CREBBP | splice | Preserved subclone |
| 1 | FPGS | missense | Acquired |
| 1 | FPGS | splice_region | Acquired |
| 1 | FPGS | UTR_3 | Acquired |
| 1 | FPGS | UTR_3 | Lost |
| 1 | ITGA9 | missense | Acquired |

| | | | |
|---|---|---|---|
| 1 | KIF21B | frameshift | Acquired |
| 2 | KIF21B | missense | Preserved |
| 4 | KRAS | missense | Acquired |
| 4 | KRAS | missense | Lost |
| 1 | KRAS | missense | Lost to subclone |
| 3 | KRAS | missense | Preserved |
| 3 | KRAS | missense | Preserved from subclone |
| 1 | KRAS | missense | Preserved subclone |
| 1 | MSH6 | missense | Acquired |
| 2 | NR3C1 | frameshift | Acquired |
| 1 | NR3C1 | splice | Acquired |
| 1 | NR3C1 | frameshift | Preserved |
| 3 | NRAS | missense | Acquired |
| 9 | NRAS | missense | Lost |
| 5 | NRAS | missense | Preserved |
| 3 | NRAS | missense | Preserved from subclone |
| 2 | NRAS | missense | Preserved subclone |
| 5 | NT5C2 | missense | Acquired |
| 1 | NT5C2 | proteinIns | Acquired |
| 1 | NT5C2 | splice | Acquired |
| 1 | NTRK3 | missense | Acquired |
| 1 | NXPH2 | UTR_3 | NA |
| 3 | PRPS1 | missense | Acquired |
| 1 | SULF2 | missense | Acquired |
| 2 | SULF2 | silent | Acquired |
| 1 | SULF2 | missense | Preserved |
